# RIFINs displayed on malaria-infected erythrocytes bind both KIR2DL1 and KIR2DS1

**DOI:** 10.1101/2024.04.30.591854

**Authors:** Akihito Sakoguchi, Samuel G. Chamberlain, Alexander M. Mørch, Thomas E. Harrison, Michael L. Dustin, Hisashi Arase, Matthew K. Higgins, Shiroh Iwanaga

## Abstract

To discriminate between our own cells and foreign cells such as a pathogen, natural killer cells are armed with inhibitory and activating immune receptors. In some cases, such as the KIRs, these are found in pairs, with inhibitory and activating receptors containing nearly identical extracellular domains that are coupled to different intracellular signalling domains^1^. The balance in signalling mediated by these receptors determines whether an NK cell is activated to destroy a target cell. Previous studies have shown that RIFINs, displayed on surfaces of *Plasmodium falciparum*-infected erythrocytes, can bind to inhibitory immune receptors and dampen NK cell activation^2,3^, reducing parasite killing. Here, we identify a clade of RIFINs that bind to inhibitory immune receptor KIR2DL1 approximately ten-times more strongly than KIR2DL1 binds its host ligand, MHC class I. We show that this interaction mediates inhibitory signalling and reduces activation of KIR2DL1-expressing NK cells. We reveal the structural basis for KIR2DL1 binding and show that the RIFIN binding surface of KIR2DL1 is conserved in the activating immune receptor KIR2DS1. We find that KIR2DL1-binding RIFINs can also bind to KIR2DS1 and that these RIFINs cause activation of KIR2DS1-expressing NK cells. This highlights the evolutionary battle between pathogen and host, suggesting that activating KIRs may have evolved to allow detection of red blood cells infected with *Plasmodium falciparum*, helping the host to clear the parasite.

## Main

The most-deadly human-infective malaria parasite, *Plasmodium falciparum* places protein molecules onto the surfaces of infected red blood cells (iRBC). These are often encoded by multigene families, of which the RIFINs are the largest. Clades of RIFINs have been identified which bind inhibitory receptors, such as leukocyte Ig-like receptor B1 (LILRB1) and leukocyte associated Ig-like receptor 1 (LAIR1). LILRB1-binding RIFINs have been shown to suppress NK cell function and are likely to reduce clearance of the parasite by host immunity^2^. A correlation between high anti-RIFIN antibody titres and less severe disease has been observed in infected children in Gabon^4^. Moreover, people living in malaria-endemic areas across Africa have unique antibodies, containing exons of LILRB1 and LAIR1, which can bind multiple RIFINs^5,6,7^. Therefore RIFIN-mediated suppression of the host immune response is likely to facilitate parasite survival and the host may adopt mechanisms to recognise RIFINs to aid protection against malaria.

Killer Immunoglobulin-like Receptors (KIRs) are expressed on NK cells and comprise pairs of inhibitory and activating immune receptors. Both inhibitory and activating KIRs possess highly similar extracellular domains. However, they have differences in their intracellular domains, resulting in opposite outcomes from signalling. There are seven types of inhibitory KIRs (KIR2DL1, 2, 3, 5 and KIR3DL1-3) and six types of activating KIRs (KIR2DS1-5, and KIR3DS1)^8^. Inhibitory KIRs play a crucial role in mediating the ‘missing-self’ theory of NK cells, in which virus-infected cells have reduced levels of the inhibitory KIR ligand, HLA class I^9^. This results in reduced KIR-mediated inhibitory signalling and in increased NK cell-mediated elimination of infected cells^10^. Activating KIRs are proposed to have evolved from inhibitory KIRs through mutations in the ITIM domain^11^. They are presumed to recognize surface-displayed molecules on pathogens, thereby activating NK cells to facilitate clearance of pathogens^12^. However, no pathogen-derived protein ligand for an activating KIRs had yet been identified, leaving their biological roles during infection uncertain.

### RIFINs from field isolates and laboratory-adapted strain of *Plasmodium falciparum* bind to inhibitory immune receptor KIR2DL1

The discovery of RIFINs which bind to inhibitory immune receptors, coupled with the structural similarity of KIRs to LILRs^13^ and LAIRs^14^, led us to ask whether we could identify RIFINs that interact with inhibitory KIRs. We first assessed whether four field-isolated strains of *P. falciparum* interact with seven fluorescently-labelled inhibitory KIRs by flow-cytometry. A small fraction (5.4%) of iRBCs infected with *P. falciparum* strain Lek174 bound to APC-labelled KIR2DL1-Fc (Fig 1a, Extended data Fig. 1). In contrast, we did not detect significant binding between other inhibitory KIRs and Lek174-infected iRBCs (Extended data Fig. 1). Neither did we detect binding of any inhibitory KIR-Fc to the remaining three parasite strains tested (Extended data Fig. 1).

**Figure 1:**
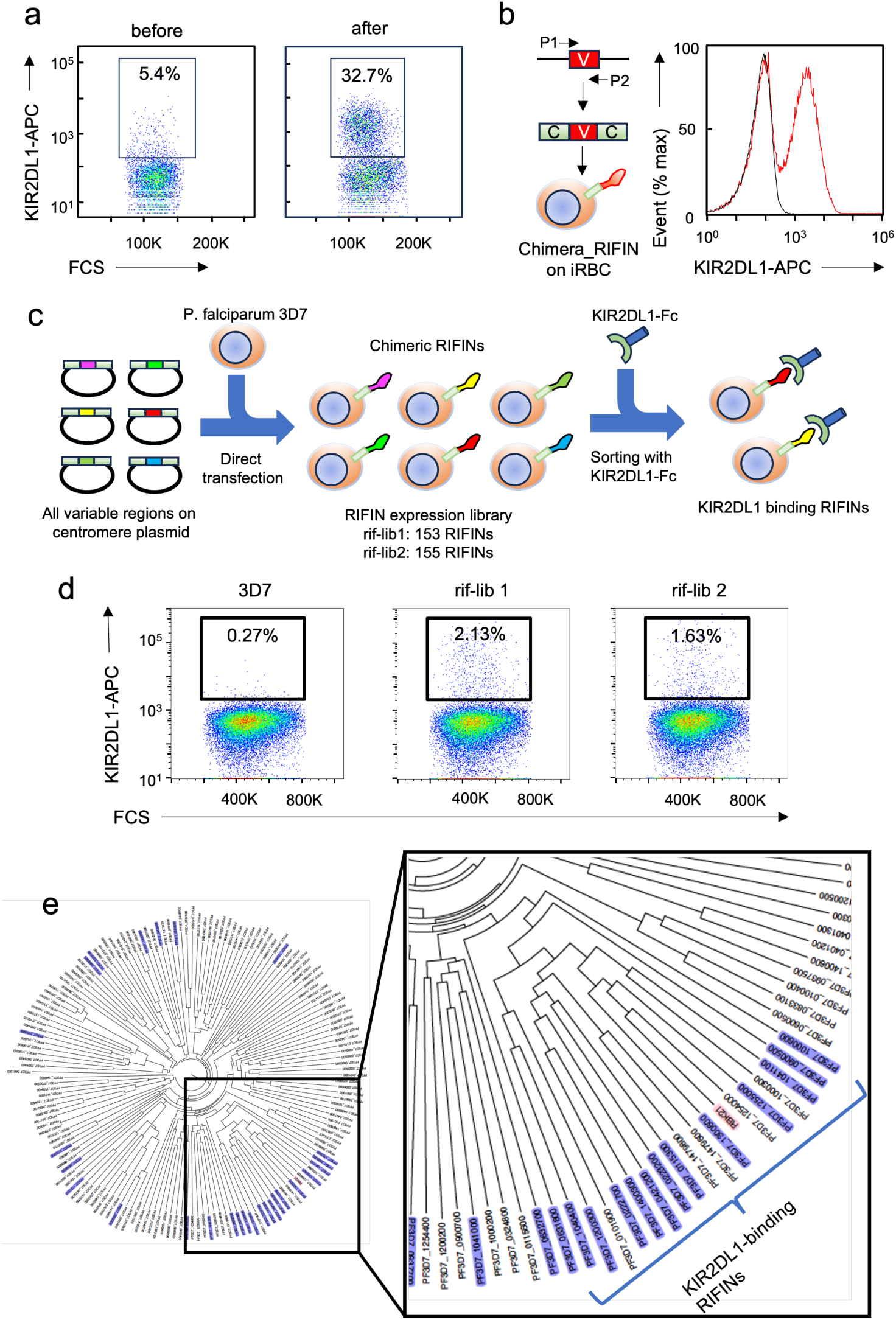
Identification of KIR2DL1-binding RIFINs. **a)** Flow sorting of lek174-infected RBCs, before (left) and after (right) selection using fluorescent KIR2DL1-Fc. **b)** A chimeric KIR2DL1-binding RIFIN containing the variable domain of RBK21 was expressed on the surface of iRBC. Flow-sorting was used to study binding of fluorescent KIR2DL1-Fc to *P. falciparum* 3D7 strain (black) and to the transgenic parasite expressing RBK21 (red). **c)** Schematic overview of generation of RIFIN expression library and its screening for binding to KIR2DL1-Fc. **d)** Flow sorting of iRBC expressing KIR2DL1-binding RIFINs from 3D7 strain or rif-lib1 and - lib2 using fluorescent KIR2DL1-Fc. These iRBC were further sorted and used for identifying KIR2DL1-binding RIFINs with NGS. **e)** Phylogenetic tree of the variable regions of RIFIN from the *P. falciparum* 3D7 strain. KIR2DL1-binding RIFINs identified from rif-lib1 and -lib2 are highlighted in violet with RBK21 in pink. The inset box shows the cluster of KIR2DL1-binding RIFINs.

We next assessed whether a RIFIN was responsible for binding Lek174-infected iRBCs to KIR2DL1. As LILRB1-, LILRB2- and LAIR1-binding RIFINs use their variable domains to bind ligands, we enriched KIR2DL1-binding cells from iRBCs infected with parental Lek174 by cell-sorting and amplified the cDNA fragments of the RIFIN variable domains (Fig. 1a). A single cDNA was specifically amplified. We next generated a transgenic parasite line which expressed a chimeric RIFIN consisting of the variable domains of this putative KIR2DL1-binding RIFIN, fused to the conserved region of a LILRB1-binding RIFIN (PF3D7_1254800) (Extended data Fig. 2a), and showed that this line bound KIR2DL1-Fc (Fig. 1b). We further cloned the full-length cDNA of this RIFIN and confirmed binding of iRBCs infected with a parasite line expressing this RIFIN to KIR2DL1-Fc fusion protein (Extended data Fig. 2b). We name this RIFIN RBK21 (RIFIN that Binds to KIR2DL1).

**Figure 2:**
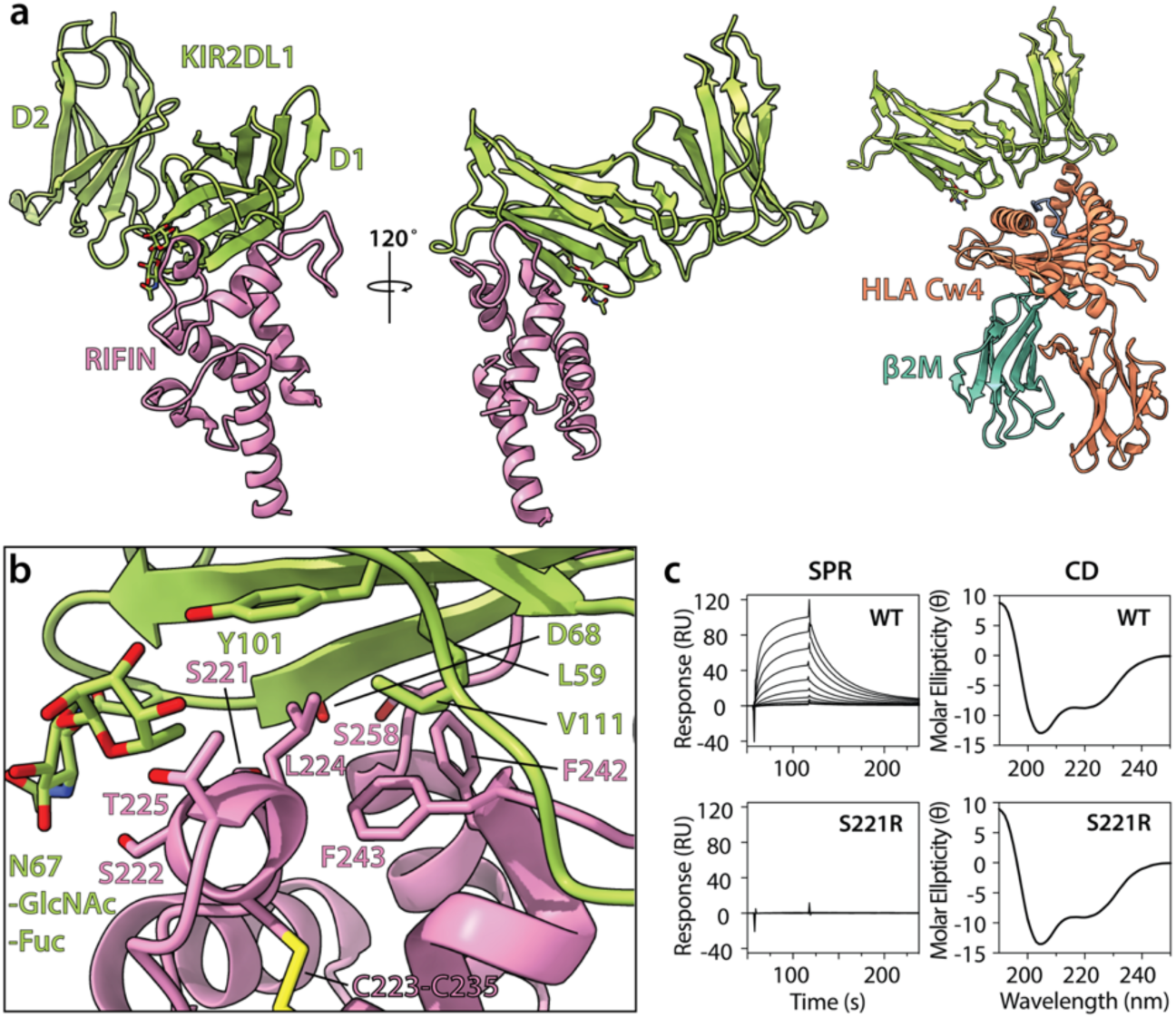
Structural basis of RIFIN binding to KIR2DL1. **a)** Two views of the complex of KIR2DL1 (green) and RBK21 (pink). The right-hand panel shows the complex of KIR2DL1 (green) bound to the MHC class I molecule, HLA Cw4 (orange and teal) (PDB: 1IM9). **b)** The interface between RBK21 and KIR2DL1 with labels for interfacial residues and for the disulphide bond which secures the binding region of the RIFIN. **c)** Experiments comparing the binding (SPR, left) and folding (CD, right) of both WT RBK21 and S221R mutant. For SPR, two-fold dilutions from 20 µM to 39 nM of either WT or S221R RBK21 were flowed over immobilised LILRB1. The same RBK21 variants were tested for correct folding by comparison using CD spectroscopy.

As the *P. falciparum* genome contains over 150 RIFINs, we next investigated whether parasites have multiple RIFINs that bind to KIR2DL1. However, as transcription of most RIFINs is usually suppressed by epigenetic control mechanisms in wild-type parasites, conducting a comprehensive analysis of RIFINs using wild-type parasites is not feasible. We therefore developed a RIFIN expression library by pooling genetically engineered parasites in which individual RIFINs were forcibly expressed, allowing analysis of the complete RIFIN repertoire (Fig. 1c). To generate this library, DNA fragments of RIFIN variable regions were amplified from 3D7 strain genomic DNA using degenerate primers which can amplify all RIFINs and were ligated into centromere plasmid, pFCEN_rif, between two conserved regions from PF3D7_1254800 (Extended Data Fig. 2a). This resulted in chimeric RIFINs containing the ligand-binding variable domains of the 3D7 RIFINs. The pool of chimeric RIFINs genes were electroporated into a recipient 3D7 strain. Library construction was carried out in duplicate, resulting in two biologically independent RIFIN expression libraries, named rif-lib1 and rif-lib2. To examine whether these contained all RIFINs, of which there are 157 in the 3D7 genome, we recovered variable regions and sequenced by amplicon-seq. This identified 150 and 153 variable regions of RIFINs in rif-lib1 and -lib2, respectively, showing that both libraries covered almost all RIFINs from strain 3D7 (Extended Data Table 1 and 2).

We next assessed binding of these libraries to KIR2DL1-Fc by flow sorting and found 2.13% of positive iRBC in rif-lib1 and 1.67% in rif-lib2 (Fig. 1d, Extended Data Table 3 and 4). Enriched iRBC libraries were then cultured, allowing us to identify KIR2DL1-binding RIFIN candidates in selected iRBCs by amplicon-seq of recovered pFCEN_rif plasmids. 19 and 24 RIFINs were identified as candidates from rif-lib1 and -lib2, respectively, with 16 of these present in both biologically independent screens (Extended Data Table 5 and 6).

To investigate the relationship between these KIR2DL1-binding RIFINs, we conducted phylogenetic analysis based on amino acid sequences of variable regions. Ten of the 16 KIR2DL1-binding candidate RIFINs were classified into the same clade of this tree, indicating that they likely evolve from a common ancestral RIFIN (Fig. 1e and Extended data Fig. 3). Interestingly, RBK21 was found in the same clade, showing that this evolutionary relationship is conserved amongst *P. falciparum* strains (Fig 1e). This clade consists of 17 RIFINs, of which 10 were identified by library screening. To determine whether the entire clade binds to KIR2DL1 we generated transgenic parasites expressing each of the 17 RIFINs and discovered that 15 of 17 RIFIN could bind to KIR2DL1-Fc. (Extended data Fig.4). The two RIFINs that did not bind to KIR2DL1 were also not identified in the library screen and may have lost KIR2DL1 binding during evolution. The conservation of KIR2DL1 binding RIFINs suggests there is a major selective advantage to targeting the KIR2DL1 inhibitory immune receptor.

**Figure 3:**
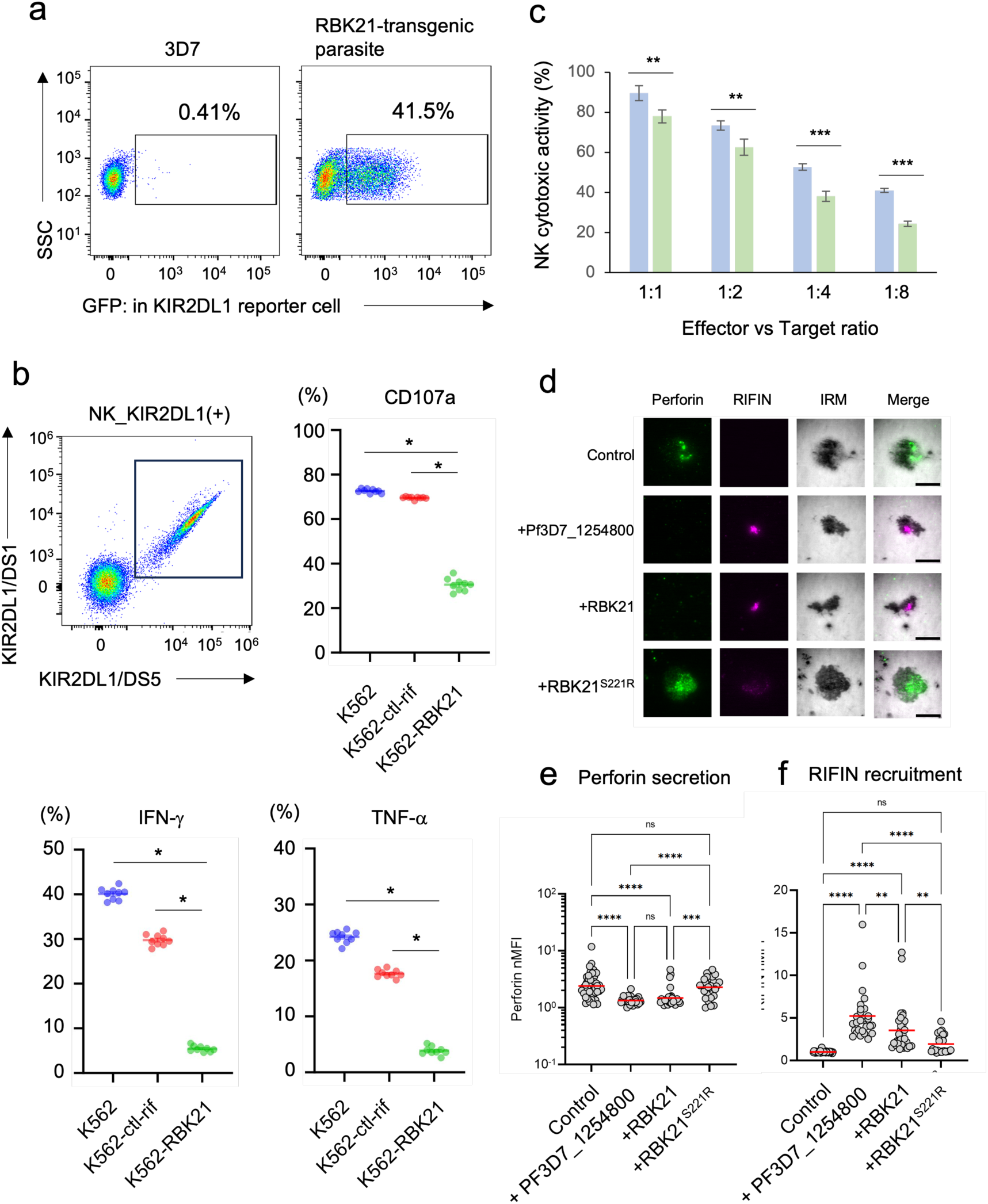
Functional analysis of RBK21. **a)** GFP expression in KIR2DL1-reporter cells upon stimulation with iRBCs infected with parental 3D7 cells (left) and an RBK21-expressing parasite line (right). Percentage of GFP-positive cells are shown. **b)** KIR2DL1-positive NK are selected using anti-KIR2DL1/DS1 and anti-KIR2DL1/DS5 antibodies (top-left panel). The double positive subset express KIR2DL1. The effect of K562 parental cells or K562 cells expressing a RIFIN which doesn’t bind KIR2DL1 (ctl-rif) or K562 cells expressing RBK21 on these KIR2DL1 expressing cells was assessed. CD107a expression (top right) and IFNg and TNFa production (bottom left and right, respectively) were each measured and the percentage of positive cells compared. Data represent the mean ± SD (n=9 technically independent samples) The original data are shown in Extended data Figure 9. *P < 0.05 (two-sided Student’s t-test). **c)** Effect of RBK21 on cytotoxic activity of KIR2DL1-NKL cell were assessed. For each target:effector ratio this was conducted using K562 cells expressing RBK21 (green) or parental K562 cells (blue) as targets. Data represent the mean ± SE (n=6 biologically independent samples). *P < 0.05 (two-sided Student’s t-test). **d).** Analysis of localisation of RIFINs (pink) or perforin (green) in the contact area for NK cells on SLBs coated with RIFIN Pf3D7_1254800, RBK21 or RBK21^S221R^. The top panel shows representative images with scale bars = 10 μm. **e)** Measurements from three independent donors were pooled and analysed to investigate the quantity of perforin in the contact area (left lower panel). Control (n = 43 cells), Pf3D7_1254800 (n = 28 cells), RBK21 (n = 31 cells) and RBK21^S221R^ (n = 29 cells). Each data point represents the measurement from one cell. A Dunn’s multiple comparisons test was performed. All revealed adjusted P values of ≤ 0.0002 except for control vs RBK21^S221R^ and Pf3D7_1254800 vs RBK21 (adjusted P value > 0.9999). **f)** Also quantified was the recruitment of RIFIN (lower right panel) for each cell, as before. Tukey’s multiple comparison test was performed and all had adjusted p value < 0.0056 except for control vs RBK21^S221R^ (adjusted P value > 0.1777). ^∗^p < 0.05, ^∗∗^p < 0.01, ^∗∗∗^p < 0.001, ^∗∗∗∗^p < 0.0001. MFI = mean fluorescence intensity.

**Figure 4.**
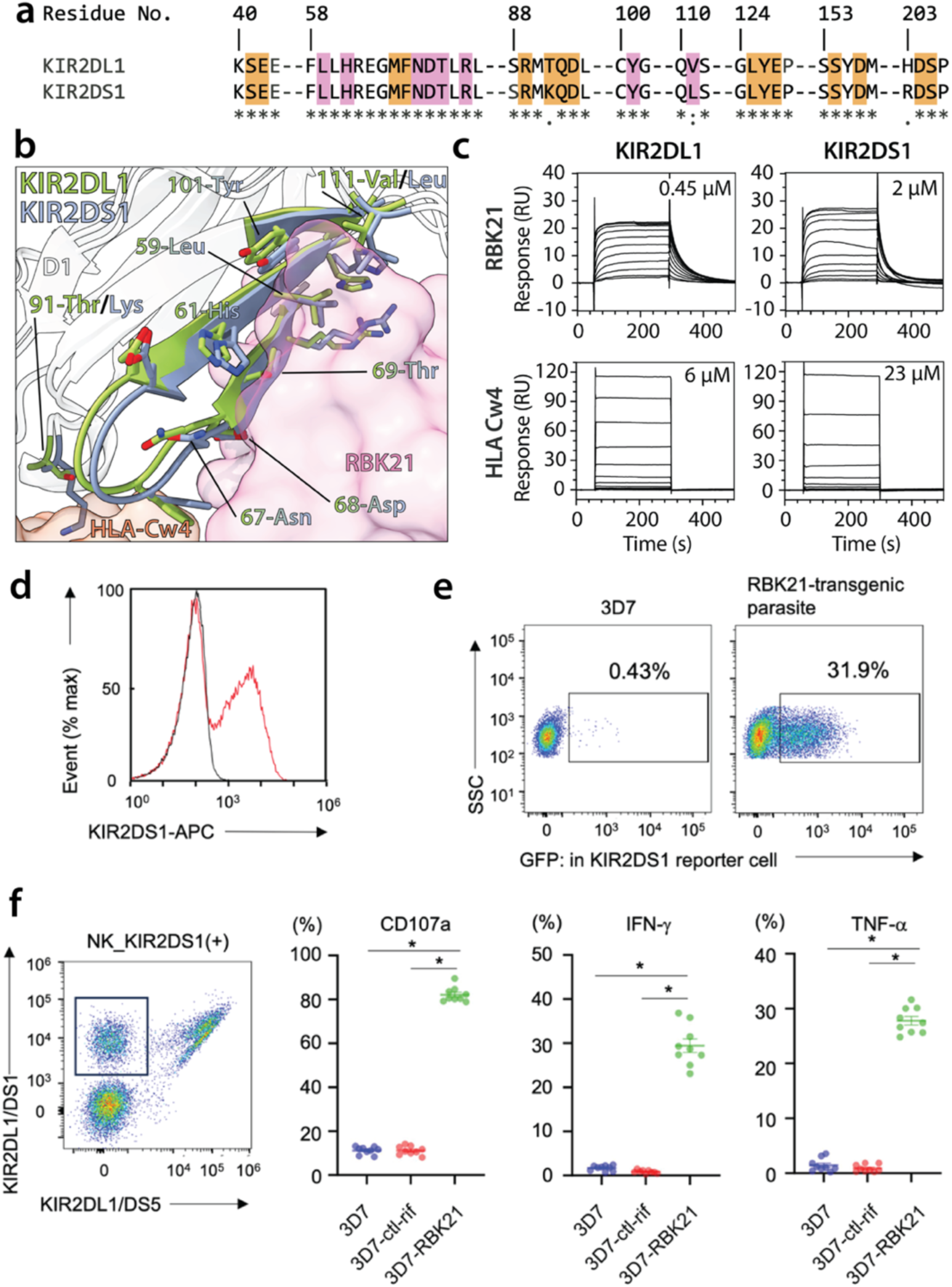
Analysis of the effect of RBK21 on KIR2DS1. **a)** Alignment of the sequences of KIR2DL1 and KIR2DS1, focused on the binding sites for RBK21 and HLA-Cw4. Residues interacting with RBK21 (pink) and HLA-Cw4 (orange) are highlighted. **b)** The structure of KIR2DL1 (green) is overlaid with a homology model of KIR2DS1 (blue). Residues which contact RBK21 (pink surface) or HLA-Cw4 (orange surface) are shown as sticks. Polymorphisms at positions 91 and 111 are highlighted. **c)** Surface plasmon resonance analysis of the binding of KIR2DL1 (left) and KIR2DS1 (right) to RBK21 (top) and HLA-Cw4 (bottom). **d)** The binding of KIR2DS1-Fc fusion protein to iRBCs infected with *P. falciparum* 3D7 (black) or with transgenic parasite expressing RBK21 (red). **e)** GFP expression in KIR2DS1-reporter cells upon stimulation with *P. falciparum* 3D7 (left) or with a transgenic parasite expressing RBK21 (right). Percentage of GFP-positive cells are shown. **f)** KIR2DS1-positive NK were selected as indicated in the box. Effect of RBK21 on CD107a expression and IFNg and TNFa production in the selected subset were assessed in the similar way as Fig. 3b. Data represent the mean ± SD (n=9 technically independent samples) The original data are shown in Extended data Figure 10. *P < 0.05 (two-sided Student’s t-test).

### The structural basis for the interaction between RIFINs and KIR2DL1

Previous structural studies have revealed how RIFINs bind to the inhibitory immune receptors LILRB1 and LAIR1^3,15,16^. These showed that RIFINs bind to the same site on LILRB1 that is bound by its host ligand, HLA class I, allowing RIFINs to signal through LILRB1 to reduce NK cell activation^2,3^. HLA class I molecules are also the ligand for KIR2DL1. To show how RBK21 binds to KIR2DL1, and to assess whether RBK21 binds to the same site on KIR2DL1 as HLA class I, we determined the structure of the KIR2DL1:RBK21 complex. An RBK21 construct comprising the variable region (residues 148-299) was mixed with the KIR2DL1 ectodomain and the complex was crystallised (Extended data Fig. 5a,b). Data were collected to 2.89 Å and molecular replacement using KIR2DL1^17^ as a search model allowed structure determination, revealing eight copies of the KIR2DL1:RBK21 complex in the asymmetric unit of the crystal (Fig.2, Extended Data Table 7).

The RIFIN adopts a mostly helical core architecture, with extended loops at the KIR-facing end. It is most similar to LILRB1-D3 binding RIFINs (Extended data. Fig. 5c)^5^ and both bind ligands using equivalent surfaces. However, these two RIFINs vary significantly in the architectures of their ligand binding loops. The KIR2DL1 binding site predominantly consists of two helices and two loops, which form extended contacts with the side of the D1 domain of KIR2DL1 (Fig. 2a,b, Extended Data Table 8). The eight copies of RBK21 in the crystal asymmetric unit only vary substantially in loop 4 (residues 244-256), which is found in two major conformations and does not contact KIR2DL1 (Extended data Fig. 6a and 6b). A comparison of the structure with that of KIR2DL1 bound to HLA class I^17^, shows that the RIFIN binding site does not overlap that of HLA class I (Fig. 2a). The contact surface between RBK21 and KIR2DL1 has an area of 878 Å^2^, with a shallow hydrophobic pocket consisting of L59, Y101 and V111 of KIR2DL1 occupied by a hydrophobic patch containing L224, F242 and F243 of RBK21. This hydrophobic core is further stabilized by an intermolecular beta sheet formed by loop 4 of RBK21, ending with a hydrogen bond formed between S258 of RBK21 and D68 of KIR2DL1. Finally, a series of polar interactions occur between S221, S222 and T225 of RIFIN helix 4 and the NAG-FUC core of a N-linked glycan on N67 of the KIR (Fig. 2b), with the glycan core clearly visible in the electron density (Extended data Fig. 5b). Within this interface lies S221 and we predicted that the S221R mutation would abolish KIR2DL1 binding. Indeed, while surface plasmon resonance (SPR) measurements showed that RBK21 bound to KIR2DL1, the S221R mutant did not bind, despite circular dichroism indicating that it adopts the correct fold (Fig 2c).

We next analysed the degree of sequence conservation among the KIR2DL1-binding RIFINs by calculating the sequence entropy for each amino acid across the known KIR2DL1-binding RIFIN clade and plotting this onto the structure (Extended data Fig. 7a). Conserved residues are found within the core of the domain, indicating maintenance of overall architecture (Extended data Fig. 7b). In contrast, the surface, except for two exposed serine residues, is highly variable in nature. This pattern of conservation of critical hydrophobic residues that maintain a core fold, coupled with high surface sequence diversity is also seen in other iRBC surface protein families, such as the DBL and CIDR⍺ domains of PfEMP1^18,19^ and the CIR proteins^20^. In a similar way, KIR2DL1-binding RIFINs use a core fold to form a binding pocket while surface diversity reduces recognition by the immune system.

### Inhibitory signalling mediated by KIR2DL1-binding RIFINs reduces cytotoxic attack of iRBCs by NK cells

The noncanonical binding of RBK21 to KIR2DL1 raised the question of whether RIFINs can transduce inhibitory signals, as observed for the endogenous ligand of KIR2DL1, HLA-class I. To test this, we used an assay in which iRBCs infected with RBK21-expressing transgenic parasites were incubated with an activated T cell reporter line (NFAT-GFP) (Fig 3a). In this reporter system, the extracellular domain of KIR2DL1 is fused to the transmembrane and intracellular domains of paired Ig-like receptor b (PILRb). On KIR2DL1 activation, this interacts with signalling adaptor, DAP12, resulting in GFP expression. Co-culture of this line with iRBCs infected with the parental 3D7 strain showed GFP expression in less than 0.41 % of the KIR2DL1-reporter cells (Fig. 3a and Extended data Fig. 8). In contrast, iRBCs infected with RBK21-expressing parasites induced strong GFP expression of 41.5 % in the reporter line (Fig. 3a).

We next investigated whether RBK21-mediated signalling suppresses NK cell activity. We examined the effect of RBK21 on NK cells function by measuring expression of CD107a, which is a marker of degranulation, as well as measuring production of the cytokines IFN-γ and TNF-α. Peripheral blood mononuclear cells (PBMC) were obtained from KIR2DL1-positive donors, and the effect of co-culturing these with K562 cells was monitored. When PBMCs were co-cultured with normal K562 cells, CD107a expression, and IFN-γ and TNF-α production were detected in 72.6±0.3%, 40.1±0.4%, and 24.2±0.4% of the KIR2DL1-positive NK cell, respectively, suggesting a substantial fraction of activated cells (Fig. 3b and Extended data Fig. 9). Similar CD107a expression and IFN-γ and TNF-α production (69.6±0.2%, 29.8±0.4%, and 17.6±0.2% of KIR2DL1-positive NK population) were detected when PBMCs were co-cultured with K562 expressing a RIFIN that does not bind to KIR2DL1 (PF3D7_1254200). In contrast, when K562 cells expressing RBK21 were cocultured with PBMCs obtained from a KIR2DL1-positive donor, significant lower CD107a expression (30.5±1.0%) and IFN-γ (5.4±0.2%) and TNF-α (3.8±0.3%) production were detected in the KIR2DL1-positive subset of NK cells, showing that RBK21 inhibits NK function through KIR2DL1-mediated signalling. Indeed, RBK21-expressing K562 cells were also able to suppress the cytotoxic activities of a KIR2DL1-positive NKL cell line in NK killing assays (Fig. 3c).

We next used a previously described supported lipid bilayer (SLB) assay^3^ to image the effect of RBK21 on NK cell activation (Fig 3d). Primary NK cells were purified from three independent donors and were incubated with SLBs which display both ICAM-1, to mediate adhesion, and PfRH5, together with a human PfRH5-binding antibody, to trigger antibody-dependent cellular cytotoxicity (ADCC). Some bilayers were also coated with RBK21, the non-binding RBK21^S221R^ mutant or the LILRB1-binding RIFIN, Pf3D7_1254800 as a positive control. Immune synapses were imaged using total internal reflection fluorescence-microscopy, allowing visualisation of RIFIN accumulation and of perforin deposition (Fig. 3d)^21^. Synapses on bilayers which lacked RIFINs showed perforin deposition, while bilayers decorated with Pf3D7_1254800 showed RIFIN accumulation and reduced perforin deposition, as previously observed^3^. When RBK21 was present in the bilayer, it was recruited to the immune synapse and resulted in a significant reduction in perforin deposition, indicating that KIR2DL1-mediated signalling had occurred. In contrast, the non-binding mutant RBK21^S221R^ was not significantly recruited to the synapse and did not inhibit perforin deposition. These data confirm that RIFIN-mediated engagement of KIR2DL1 results in inhibitory signalling that supresses NK cell activation.

### RBK21 also signals through activating immune receptor KIR2DS1 and increases the cytotoxic function of KIR2DS1-expressing NK cells

The KIR family contains both inhibitory and activating receptors, which have largely conserved extracellular domains. Inspection of the RBK21-KIR2DL1 structure revealed that the KIR2DL1 residues which mediated the interaction with RBK21 interface are almost completely conserved in KIR2DS1 (Fig. 4a and b), except for Val111, which is leucine in KIR2DS1. In contrast, a change in position 91 (Thr in KIR2DL1 and Lys in KIR2DS1) alters the MHC class I binding site (Extended Data Figure 10). We therefore used SPR analysis to quantify the binding of KIR2DL1 and KIR2DS1 to RBK21 and to the MHC class I molecule, HLA_Cw4. This shows that KIR2DS1 binds with a 2 μM affinity to RBK21, close to the 0.45 µM affinity of the RBK21-KIR2DL1 complex, while its affinity for HLA_Cw4 is over 10-fold weaker (Fig. 4c). Similarly, FACS analysis demonstrated that transgenic parasites expressing RBK21 bind to KIR2DS1 as well as to KIR2DL1 (Fig. 4d). These findings suggest that KIR-binding RIFINs are novel pathogenic ligands for activating KIR receptor.

Using the NFAT-GFP reporter system with KIR2DS1, we next examined whether binding of RBK21 to KIR2DS1 resulted in signal transduction (Fig 4e). Robust expression of GFP was found in KIR2DS1-expressing reporter cells co-cultured with RBK21-expressing parasites (31.9%), compared to the control (0.4%). We also once again assessed the effect of iRBCs infected with RBK21-expressing transgenic parasites on KIR2DS1-positive NK cells by monitoring the markers CD107a, TNF⍺ and IFNγ in co-culture. This revealed an RBK21-dependent stimulation of CD107a expression (82.1±1.1%) and of IFN-γ (29.4±1.5%) and TNF-α (27.8±0.8%) production, which was not observed when using iRBC infected with parasites expressing the negative control RIFIN (Fig. 4f and Extended data Fig. 11). Taken together, these data show that as well as this RIFIN engaging the inhibitory KIR2DL1 to suppress the immune response, it can also bind to activating receptor KIR2DS1. However, KIR2DS1 binding has the opposite effect of activating NK cells by triggering both cytotoxic and cytokine responses. As different NK cells express different KIRs, this will equip our immune systems with the ability to clear iRBCs expressing KIRDS1-binding RIFINs.

## Discussion

The importance of activating immune receptors in mice was demonstrated in Murine Cytomegalovirus (MCMV) infection^22^. Mouse NK cells possess inhibitory and activating C-type lectin Ly49 receptors, equivalent to KIRs in humans^23^. Inhibitory Ly49 receptors, such as Ly49I, recognize MHC-class I molecules as endogenous ligands and regulate NK cell function through the “Missing Self theory”, in an equivalent way to inhibitory KIRs^24^. In contrast, activating Ly49 receptor, Ly49H, binds to the MCMV-derived molecule m157, which is expressed on infected cell surfaces^22^ triggering cytotoxic activity of NK cells. Our results demonstrate that, as observed in mice, humans also use activating receptors to actively engage NK cells in defence against infection. In both mice and humans, although the identity of the activating receptors differ, hosts have evolved inhibitory receptors into activating receptors in response to pathogens that acquire ligand molecules for those inhibitory receptors.

ADCC is an adaptive immune response largely mediated by NK cells, in which the CD16 (FCγRIII) receptor recognizes the Fc portion of pathogen-interacting IgG antibodies^25^. *In vitro* studies have recently highlighted the ability of NK cells to destroy malaria parasite-infected RBCs, with recognition mediated by antibodies targeting iRBC surface proteins^26^. Recent epidemiological studies conducted in Ugandan children have shown that repeated infections with *P. falciparum* alter the composition of NK cell populations, resulting in an increase in the ratio of CD56neg NK cells^27^. These CD56neg NK cells represent an atypical subset of NK cells distinct from the typical CD56dim NK cells and exhibit stronger ADCC activity compared to CD56dim NK cells, perhaps improving NK cell mediated iRBC clearance. In response, parasites may evade ADCC by using RIFINs to stimulate inhibitory immune receptors, such as LAIR1, LILRB1 and KIR2DL1, thereby inhibiting the activation of NK cells on recognition of iRBCs through CD16.

Paired receptors evolved under a selection pressure that resulted in retention of extracellular homology between the members of the activating/inhibitory pair. This is true in the case of KIR2DL1/S1, with the RIFIN-binding interface remaining almost perfectly conserved between the pair. In contrast, at the MHC-binding site, KIR2DS1 contains a threonine to lysine mutation that reduces binding to the MHC class I molecule, HLA-Cw4. This will contribute to the inability of KIR2DS1-expressing NK cells to produce activating signals when co-cultured with cells expressing conventional HLA class-I molecules, thereby avoiding targeting of host cells^28^. Our findings therefore suggest that the evolutionary origin of activating KIRs is a response to *Plasmodium falciparum* successfully co-opting the immune response through inhibitory KIRs, with activating receptors evolving point mutations which retain RIFIN binding in the low micromolar range, while reducing binding to its canonical ligand, MHC class I. Indeed, previous work has established that low KIR2DS1 expression is common in regions with historically low malaria transmission, where it will not be required for malaria prevention^29^.

While the host generates genetic diversity through paired receptors, the parasite has been shaped by the evolutionary pressure to avoid immune detection, with antibodies to RIFINs commonly found in natural malaria infections^4,5,30,31^. This is highlighted when we map sequence diversity within the KIR2DL1-binding clade onto the structure of the RIFIN. This clade of RIFINs conserves a minimal motif of core hydrophobic residues which maintain the basic scaffold necessary to target KIR2DL1, while almost all surface residues, including those at the interface, are highly variable, most likely generating antigenically distinct molecules. Despite this low level of sequence conservation, phylogenetic analysis did allow for identification of KIR2DL1 binding RIFINs missed in the initial screen and might provide an important tool for identification of future RIFIN families and which immune receptors they target. Importantly, broadly reactive antibodies that target multiple RIFINs through inclusion of whole domains of LILRB1 and LAIR1 have been discovered^5^, providing the host with antibodies that allow detection of this large protein family.

In summary, through identification of a RIFIN that targets both KIR2DL1 and KIR2DS1, we have revealed more about the evolutionary battle between malaria parasite and host. The acquisition of KIR-binding RIFINs likely allowed the parasite to suppress NK cell function in the presence of ADCC. In response, an offensive strategy using activating KIR allows the host to target the same RIFINs that engage inhibitory immune receptors to instead trigger activating immune signalling, potentially equipping the host with the ability to overcome parasite immune evasion. The implication of this evolutionary arms race warrants further investigation, with more focussed epidemiological study of KIR diversity across malaria-endemic regions, in addition to studies investigating the feasibility of targeting RIFINs in future therapeutic interventions.

## Materials and Methods

### Ethical Statement

Erythrocytes and human serum were obtained from the Japanese Red Cross (research ID:25J0143) with written informed consent. Parasites had been collected from Thai patients according to the ethical approvals by Chiang Mai University (permission number: 187/2554) and Mie University (Permission number: 1312), and their use was approved by the University of Osaka (permission number: 149-003).

### Cells and antibodies

NKL cells were generously provided by L. L. Lanier at the University of California, San Francisco. The human erythroleukemia cell line, K562, was sourced from the Cell Resource Centre for Biomedical Research, Institute of Development, Ageing and Cancer, Tohoku University. Anti-KIR2DL1/DS1 (Milteny Biotec, 130-118-973) and anti-KIR2DL1/DS5 (R&D system, MAB1844-SP) antibodies were purchased and used for detection of KIR2DL1(+)- and KIR2DLS1(+)-NK cells in PBMC. The PBMC were obtained from healthy donors. Anti-FLAG antibody (Sigma-Aldrich, F1804) and allophycocyanin (APC)-conjugated anti-human IgG Fc antibody (Jackson ImmunoResearch,109-136-098) were used for detection of transgenic K562 and for binding assay involving Fc-fusion proteins, respectively. PacificBlue-conjugated anti-human CD107a (Biolegend, 328623), FITC-conjugated CD56 (Biolegend, 318303), PerCP/Cy5.5-conjugated anti-human IFNg (Biolegend, 506527), and APC/Cy7-conjugated anti-human TNFa (Biolegend, 502943) antibodies were purchased and used for functional assay of RBK21.

### Parasite culture

1. *P. falciparum* was cultured with human RBCs (type O blood, hematocrit 2%) in complete medium (CM). This medium consisted of RPMI-1640 medium containing 10% human serum and 10% AlbuMAXI (Invitrogen), along with 25 mM HEPES, 0.225% sodium bicarbonate, and 0.38 mM hypoxanthine. Additionally, the medium was supplemented with 10 mg/ml gentamicin. Cultures were maintained under an atmosphere containing 90% N_2_, 5% CO_2_, and 5% O_2_. Parasites were routinely synchronized every 4 days using a 5% sorbitol solution and subsequently utilized for analyses.

### Cloning of RBK21 cDNA

The iRBCs that bound to KIR2DL1-Fc were enriched using the SH800 cell sorter (SONY). The detailed procedure for sorting iRBCs follows. Enriched iRBCs were cultured, and total RNA was purified using TRIzol (Thermo Fisher Scientific). Subsequently, cDNA was synthesized using SuperScriptIII reverse transcriptase (Thermo Fisher Scientific) and random hexamers according to the manufacturer’s instructions. The variable regions of RIFIN expressed in enriched iRBCs were amplified using primers shown in Extended table 9. The obtained fragment, which encoded a part of RBK21 gene, was cloned into the centromere plasmid^32^, pFCEN-rif, resulting in a chimeric RIFIN. Expression was controlled by the promoter of elongation factor a of *P. berghei*. Prior to cloning full-length RBK21 cDNA, 5’ and 3’-ends of RBK21 cDNA were analyzed using 5’-Full and 3’-Full RACE core sets (TAKARA) with specific primers (Extended Data Table 8). Following the sequencing of the amplified product, a DNA fragment encompassing the entire coding region of RBK21 was obtained through PCR. The amplified fragment was cloned in the pFCEN-rif and used for further study.

### Transfection of P. falciparum

Transfection of *P. falciparum* was carried out as previously described^33^. In brief, schizonts of *P. falciparum* 3D7 strain were purified using a percoll/sorbitol gradient solution, followed by culturing with fresh RBCs for 4 hours. Subsequently, parasites were synchronized to within a 4-hour window by treatment with 5% sorbitol. Full mature schizonts were purified from highly synchronized parasites and then transfected with pFCEN-rif plasmids with a Amaxa nucleofector 2 with parasite nucleofector kit 2 (LONZA). Since the pFCEN-rif has the human dihydrofolate reductase as a drug selectable marker, the transgenic parasites could be screeded by drug treatment with pyrimethamine. Transfection experiments using each plasmid were carried out in duplicate throughout this study, resulting in biological independent transgenic parasites.

### Flow cytometry

The binding of KIRs-Fc fusion proteins to iRBCs was assessed by flow cytometric analysis with Attune NxT (ThermoFisher Scientific). Prior to the assay, the KIRs-Fc fusion proteins were mixed with an allophycocyanin (APC)-conjugated anti-human IgG Fc antibody. The iRBCs were mixed with the APC-labeled KIRs-Fc binding proteins, and parasite nuclei were stained with Hoechst 33342. The iRBC population were gated based on fluorescence from Hoechst 33342 and KIR-Fc bound iRBCs were selected using APC fluorescence (Supplementary Gating Strategy for Flow Cytometry Analysis 1). All obtained data were analyzed using FlowJo software (Becton Dickinson).

The iRBCs binding to KIR2DL1 were selectively sorted by the cell sorter, SH800 (SONY). Prior to sorting, the iRBC containing late trophozoite and schizonts were separated form uninfected RBC using percoll/sorbitol gradient solution, followed by mixing with APC-labelled KIR2DL1-Fc fusion protein. After binding KIR2DL1-Fc to iRBC, they were suspended in CM medium and then subjected to SH800. The sorted iRBC was immediately recovered in CM medium and cultured with fresh RBCs.

### Preparation of RIFIN expression library

DNA fragments which encoded all RIFIN variable regions of were amplified from genomic DNA of the 3D7 strain using degenerated primers, designed at the internal sites of N- and C-terminal conserved regions of RIFINs. The amplified fragments representing variable regions of all RIFINs of 3D7 strain were cloned into the *Bsm*BI sites of the pFCEN-rif plasmid (Extended Fig. 2a). The cloned fragments are flanked with parts of N- and C-terminal conserved regions on the pFCEN-rif, resulting in chimeric RIFINs genes. The pFCEN-rif plasmids containing chimeric RIFIN genes were introduced into the *P. falciparum* 3D7 strain via a single electroporation. Transgenic parasites were subsequently obtained through treatment with pyrimethamine, resulting in the creation of an RIFIN expression library. Each individual parasite within the RIFIN expression library expresses a distinct chimeric RIFINs. Library construction was carried out twice, resulting in establishment of two biologically independent RIFIN expression libraries, designated rif-lib1 and rif-lib2.

To identify selected RIFINs, the pFCEN-rif plasmids containing chimeric RIFIN genes were recovered from rif-lib1 and -lib2. The variable regions integrated into the plasmids were subsequently amplified through PCR using primers designed based on the plasmid backbone sequence. Following this, sequence tags were introduced at the ends of the amplified DNA fragments through PCR, and the resultant tagged DNA fragments were then sequenced using Miseq (Illumina). The reads containing the variable regions were isolated from the obtained fastq files using the seqkit program^34^ with the “grep” option. Subsequently, the plasmid backbone sequence was trimmed using trimomatic-0.39-2 program^35^. After confirming the sequence quality of trimmed reads, they were mapped onto the reference genome sequence of *P. falciparum* 3D7 strain, which was downloaded from PlamoDB 62, using bowtie2 tool^36^. Reads aligned to more than two different genomic loci were eliminated from the mapping data. The number of reads for each RIFIN gene was tallied using featureCounts^37^ and normalized by dividing by the total number of mapped reads.

### Screening for KIR binding RIFINs from the RIFIN expression libraries

The iRBCs expressing the KIR2DL1-biding RIFIN library were incubated with KIR2DL1-Fc fusion protein and screened using a cell sorter. They were cultured in CM immediately after sorting. The pFCEN-rif containing candidate genes of KIR2DL1-binding RIFNs were recovered from those sorted iRBCs. The DNA fragments of variable regions were amplified through PCR and tagged in the similar manner as described in the previous section. The resultant tagged fragments were sequenced using Miseq (Illunina) and the number of mapped reads were tallied and normalized. After screening, the normalized read count for each RIFIN gene was divided by the normalized read count obtained before screening. Candidates for KIR2DL1-binding RIFINs were identified based on values exceeding 2. Each variable region of identified candidates were amplified from the genomic DNA of P. falciparum 3D7 strain and individually cloned into pFCEN-rif. Transgenic parasites that have the pFCEN-rif containing those candidates were generated and their binding to KIR2DL1 was confirmed using Attune NxT and Fc fusion protein.

### Phylogenetic analysis of RIFINs

Amino acid sequences of all RIFINs of *P. falciparum* 3D7 strain were obtained from PlasmoDB (https://plasmodb.org/plasmo/app/) and aligned using Clustal omega (https://www.ebi.ac.uk/jdispatcher /msa/clustalo). Since the variable region of LILRB1-binding RIFN (PF3D7_1254800) had already been determined^3^, the corresponding regions were designated as the variable regions of each RIFIN. These obtained variable regions were aligned again using Clustal Omega, followed by generating phylogenetic tree based on newly aligned sequences. All analyses were performed with default setting. The resultant data of phylogenetic analysis was visualized using treeview program^38^.

### Transfection of mammalian cells

Stable transfectants of K562-RBK21 and NKL-KIR2DL1 were generated using retrovirus-mediated transduction with the pMxs retroviral expression vector and PLAT-E retroviral packaging cells transfected with amphotropic envelope as described previously^2,39^ (Cell Biolabs inc.). In brief, the variable region of RBK21 was fused with the transmembrane and cytoplasmic regions of PILRa (amino acid residue 196-256) and cloned into the pMxs plasmid. The resultant plasmid containing the fusion gene of RBK21 and PILRa was introduced into PLAT-E cells using TransIT-293 (Mirus) and recombinant retrovirus were harvested from supernatant 3 days after transfection. The full-length cDNA of KIR2DL1 was chemically synthesized (Genescript) and subsequently cloned into the pMxs plasmid. Recombinant retrovirus was then produced using a similar method as described above. The target cells, i.e. K562 and NKL, were seeded in 24 well culture plate and infected with produced recombinant virus, resulting in the K562-RBK21 and NKL-KIR2DL1.

### Reporter assay

The extracellular domains of KIR2DL1 and KIR2DS1 were fused with paired immunoglobulin-like receptor b (PILRb) and expressed on the surface of mouse T cell hybridomas that were stably transfected with NFAT-GFP and FLAG-tagged DAP12, as described previously^40^. The transmembrane domain of PILRb can transduce the signal via the DAP12 adaptor molecule in the reporter cell, resulting in the expression of GFP. The reporter cell lines, which expressed fusion proteins of KIR2DL1 and KIR2DS1 with PILRb, were stimulated by iRBC which was infected with parasite expressing RBK21 for 16 hours, and the GFP expression in them were monitored using Attune NxT, The *P. falciparum* 3D7 strain was used as negative control.

### Measurement of CD107a, IFNg, and TNF-a in NK cells

NK cells were purified using an NK cell isolation kit according to the manufacturer’s instruction from PBMCs obtained from KIR2DL1- and KIR2DS1-positive donors. Following purification, the NK cells were cultured in NK cell growth medium, which consisted of RPMI1640 supplemented with 15% FCS (GIBCO), 5% human serum, 1x MEM-NEAA, 1 mM sodium pyruvate, 100 units/ml penicillin, 100 μg/ml streptomycin, 500 U/ml human IL-2 (hIL-2), 5 ng/ml hIL-15, 10 ng/ml hIL-12, 40 ng/ml hIL-18, and 20 ng/ml hIL-21. To evaluate the inhibitory effect of RBK21 on NK cell function, purified NK cells (1.0 × 10^5^ cells) containing KIR2DL1-positive NK were co-cultured with K562-RBK21 cells (1.0 × 10^5^ cells) in a 96-well plate at 37°C for 4 hours. PacificBlue-conjugated anti-human CD107a antibody was added to the medium at the beginning of co-culturing for the measurement of CD107a expression. After 1 hour of co-culture, the NK cells were treated with BD GolgiStop reagent, which is included in the BD Cytofix/CytopermTM Plus Fixation/Permeabilization kit containing monensin (BD). The NK cells were collected by centrifugation and dead cells were stained at RT for 30 min with LIVE/DEADTM Fixable Yellow Dead cells stain kit (Invitrogen). NK cells were further stained on ice for 30 min with FITC-conjugated anti-CD56 antibody, PE-labelled anti-KIR2DL1/DS1 antibody, and APC-labelled KIR2DL1/DS5 antibody. The cells were fixed and permeabilized at 4 °C for 20 min with the fixation/permeabilization solution, which is also included in the kit. The permeabilized NK cells were then stained at 4 °C for 30 min in the dark with PerCP/Cy5.5-conjugated anti-human IFNg antibody and APC/Cy7-conjugated anti-human TNFa antibody using the BD Perm/Wash buffer. The CD107a expression, and IFN-γ and TNF-α production were assessed using Attune NxT. The K562 and K562-control_RIFIN(PF3D7_ PF3D7_1254200) were used as negative controls. To assess the activation of NK cells via KIR2DS1 by RBK21, NK cells, including KIR2DS1-positive NK cells, were co-cultured with iRBCs containing RBK21-expressing transgenic parasites. In this assay, the P. falciparum 3D7 strain and transgenic parasites expressing a control RIFIN were used as negative controls. CD107a expression, as well as IFN-γ and TNF-α production, were assessed as described above.

### Cytotoxicity assay

The suppression of the cytotoxic activity of NKL cells by RBK21 were assessed as previously described^2^. In brief, the K562-RBK21, which stably expressed RBK21-PILRa fusion, was labelled with 15 mM Calcein AM in assay medium (RPMI-1640 without phenol red, supplemented with 1% FCS) for 30 min at 37 °C, followed by washing twice with the assay medium. The NKL and NKL-KIR2DL1 cells were also washed twice with assay medium and mixed with K562 or K562-RBK21 in a 96 well plate with effector (NKL or NKL-KIR2DL1) : target (K562 or K562-RBK21) ratios ranging from 1:1 to 1:8 in triplicate. These cells were centrifuged at 250 g for 5 min and then co-cultured for 4 hours at 37 °C. To detect the spontaneous and maximal release of calcein-AM from K562 or K562-RBK21, cells were dispensed without NKL or NKL-KIR2DL1 in the plate, followed by lysing with 1 % Triton X-100 30 min before the end of the co-incubation. After co-incubation, cells were centrifuged at 1,500 rpm for 2 min, and the fluorescence of released calcein-AM were measured. Specific cytotoxicity (C, %) was calculated using the following formular: C = 100’ (mean fluorescence in coculture - mean fluorescence in spontaneous lysis)/(mean fluorescence in maximal lysis – mean fluorescence in spontaneous lysis).

### Protein Expression and Purification

The coding regions of KIR receptors, KIR2DL1*00302 (Gene ID; 3802, amino acid residues 22-245), KIR2DS1*001 (Gene ID; 3806, amino acid residues 22-245), KIR2DL2*001 (Gene ID; 3803, amino acid residues 22-245), KIR2DL3*001 (Gene ID; 3804, amino acid residues 22-245), KIR2DL5*001 (UniProt ID; A0A1B5B9Z9, amino acid residues 22-239), KIR3DL1*001 (Gene ID; 3811, amino acid residues 20-339), KIR3DL2*001 (Gene ID; 3812, amino acid residues 22-341), and KIR3DL3*001 (UniProt ID; Q8N743, amino acid residues 25-320), were cloned into the vector pCAGSS. The resultant plasmids were introduced into HEK293T cell using TransIT293 (Mirus), and these KIRs were expressed as secreted human-Fc fusion proteins. These KIRs-Fc fusion proteins were obtained from supernatants of transfected HEK293 cells.

To produce protein for structural biology, the coding sequence for KIR2DL1 (residues 27-221) was cloned into the vector pHLsec, including a C-terminal His_6_-tag and transfected into mycoplasma screened Expi293F™ GnTI^-^ cells using ExpiFectamine 293 (ThermoFisher Scientific). Mutagenesis of RBK21 constructs was carried out using the Q5 Site Directed Mutagenesis kit (New England Biolabs) following the manufacturer’s instructions. 5 days after transfection, supernatant was harvested by centrifugation at 5,000g to pellet cells, before filtering with a bottle-top 0.45 µM filter. The pH was adjusted to pH 7.5 through addition of Tris to a final concentration of 50 mM before passing over Ni Sepharose® Excel resin (Cytiva). Captured KIR2DL1 protein was washed with 50 mM Tris pH 7.5, 150 mM NaCl, 20 mM imidazole before elution with 50 mM Tris pH 7.5, 150 mM NaCl, 500 mM imidazole. Protein was incubated with EndoH_F_ overnight at 37 °C to cleave glycans.

The RIFIN variable domain construct (RBK21, residues 148-299) was cloned into a modified pENTR4LP vector including a cleavable N-terminal monomeric Fc domain tag and a C-terminal C-tag before transfection as into Expi293F™ GnTI^-^ cells as above. Supernatant was processed as above and flowed over CaptureSelect™ C-tagXL Affinity Matrix (ThermoFisher Scientific) to isolate the C-tagged protein. Captured RBK21-mFc was washed with 50 mM Tris pH 7.5, 150 mM NaCl and eluted with 2M MgCl_2_, 20 mM Tris pH 7.5 before buffer exchange into 50 mM Tris pH 7.5, 150 mM NaCl. Protein was incubated with EndoHF and TEV protease overnight at 37 °C to cleave glycans while simultaneously releasing the variable domain from the mFc tag. The Rifin was flowed over Pierce™ Protein G Agarose to remove the mFc fusion tag and the flow through collected before further purification by size-exclusion chromatography using a Superdex 75 10/300 column (Cytiva).

The HLA-Cw4-β2m complex used in the SPR experiments was produced using a refolding protocol as described previously^41^. In brief, the HLA (accession number; MH254935) and β-2-microgloblin (β2M; accession number; NM_004048) were expressed separately in *E. coli* BL21(DE3) pLysS and purified as inclusion bodies. These were solubilized in buffer containing 6M guanidine, 100 mM Tris-HCl, pH8.0, 2 mM EDTA, 0.1 mM DTT. Solubilized HLA and β2m were diluted to 10 mg/mL using solubilization buffer with the addition of 1 M DTT to a final concentration of 10 mM, and the solutions incubated at room temperature for 1 hour. Refolding buffer (100mM Tris pH8.0, 400mM L-Arg, 2mM EDTA, 3.73mM cystamine, 6.73mM cysteamine) was chilled to 4 °C before β2m was refolded by rapid dilution, to a final concentration of 2 µM. Refolding is allowed to proceed for 2 hours before addition of 10 mg/L of peptide (QYDDAVYKL) and then HLA C was added dropwise to a final concentration of 2 µM. The heterotrimer was allowed to refold for 72 hours at 4 degrees before size exclusion chromatography (S75 HiLoad 26/60) in 20 mM Tris-HCl pH 8, 100 mM NaCl.

### Crystallization, data collection and structure determination

KIR2DL1 and RBK21 were combined to a 1:1 molar ratio and the resulting complex was purified using a Superdex 75 10/300 column (Cytiva) in 20 mM HEPES, 150 mM NaCl. Initial crystallization trials were carried out using vapour diffusion in sitting drops by mixing 100 nL of protein solution (10 mg/mL) with 100 nL of well solution using commercial screens. Initial hits were obtained after 10 days at 18 °C in condition H12 (0.1 M Tris-Bicine pH 8.5, 0.1M Amino acid mix, 37.5 % v/v precipitant mix 4) of the Morpheus HT-96 screen (Molecular Dimensions). Using these initial crystals as seeds, an optimization screen was set up using the Morpheus stock solutions, screening a pH range from 8.0 to 9.0 and a final precipitant concentration between 21% and 28% by mixing 100 nL of protein solution with 100 nL of well solution and 25 nL of seed stock. The best crystals were obtained after 12 days with a well solution of 0.1 M Tris-Bicine, 0.1 M amino acid mix, 23% precipitant mix 4 and were harvested then cryo-cooled for data collection in liquid nitrogen. Data were collected on beamline ID30A-3 at ESRF at a wavelength of 0.97625 Å and then indexed using DIALS and scaled using AIMLESS resulting in a full dataset with a final resolution of 2.89 Å. The structure was solved using molecular replacement using KIR2DL1 (PDB accession code 1IM9) as a search model using Phaser MR (v.2.8.3)^42^. Building and refinement cycles were carried out using COOT (v.0.8.9.2)^43^ and BUSTER (v.2.10)^44^.

### Surface Plasmon Resonance

All experiments were carried out on a Biacore T200 instrument (Cytiva) using a running buffer of 20 mM HEPES pH 7.5, 300 mM NaCl, 0.005% v/v TWEEN-20. Proteins were desalted into this buffer using PD-10 columns (Cytiva). Sensorgrams were double referenced by subtraction of the response measured from a blank flow path, where no protein was immobilised in addition to subtraction of the response due to buffer from the protein flow path. Kinetic values were obtained using the BIAevaluation software (Cytiva), fitting data to a global 1:1 interaction model, allowing for determination of association rate constant (k_on_), dissociation rate constant (k_off_), and affinity (K_d_). Equilibrium fits of multicycle experiments were obtained using a 1:1 interaction model in BIAevaluation (Cytiva). All kinetic and equilibrium fits are contained within the source data file.

For experiments comparing its binding affinity for KIR2DL1 and KIR2DS1, RBK21 was coupled to a CM5 sensor (Cytiva) via amine chemistry to ∼150 RU. Two-fold dilution series of KIR2DL1 and KIR2DS1 were injected over the chip for 240 s with a flow rate of 30 µL/min, followed by 300 s dissociation time and regeneration between cycles with 10 mM Glycine pH 2.5 for 10 s. Affinity values were derived from both equilibrium and kinetic fits to the data.

To measure HLA_Cw4 binding, ∼300RU of each Fc-tagged KIR2DL1 and KIR2DS1 were captured via Protein A/G (ThermoFisher Scientific) pre-immobilized on a CM5 sensor (∼1500 RU on each flow cell). The same association and dissociation times as above were used. Regeneration of the surface was carried out with 10 mM Glycine pH 2.5 before more KIR2DL1 and DS1 Fc-tagged protein were re-immobilized on the chip and the next cycle was started. Affinity values for this data could only be fit using equilibrium measurements due to the speed of the kinetics making kinetic fits unreliable.

Comparison of the binding of WT and S221R RBK21 to KIR2DL1 was achieved through immobilisation of ∼300 RU of KIR2DL1 onto the surface of a CM5 sensor followed by injection of a two-fold dilution series of RBK21 variants starting from 20 µM over the sensor surface at a flow rate of 30 uL/min. Association and dissociation times were 60 and 120s respectively, with regeneration with 10 mM Glycine pH 2.5 for 10 s between cycles.

### Circular Dichroism

Proteins were desalted into 10 mM sodium phosphate pH 7.5, 100 mM NaF using PD-10 columns (Cytiva). Measurements were made on a Jasco J815 CD spectrophotometer. Experiments were recorded at 20 °C between 190 nm and 260 nm at 0.5 nm intervals with a protein concentration of 0.1 mg/mL using a 1 mm pathlength cuvette (Hellma Macro Cell 110-QS). Three equivalent protein spectra were recorded and averaged after subtraction of a buffer-only blank measurement. Data were processed using the CAPITO online webserver^45^.

### Shannon-Entropy Calculation

The variable region from each KIR2DL1-binding RIFIN identified in the clade was used to generate a multiple sequence alignment (MSA) using MUSCLE ^46^. The MSA was then used with the Protein Variability Server. Per-residue entropy values were then binned into three categories: absolute conservation, high conservation, and medium conservation with entropy values of 0, 0 – 0.5 and 0.5 – 0.8, respectively. These three bins were mapped onto the structure in Pymol.

### Supported lipid bilayer Assay

Supported lipid bilayer experiments were performed as previously described^3^. Briefly, 1,2-dioleoyl-sn-glycero-3-phosphocholine (DOPC, Avanti Polar Lipids) micelles were supplemented with 12.5% 1,2-dioleoyl-sn-glycero-3-[(N-(5-amino-1-carboxypentyl) iminodiacetic acid) succinyl]-Ni (Avanti Polar Lipids) and infused into plasma-cleaned glass coverslips affixed within a six-lane, adhesive chamber (Ibidi) for 20 minutes. Supported lipid bilayers were washed three times with HEPES buffered saline + 0.1% BSA + 1 mM CaCl_2_ + 2 mM MgCl_2_ and blocked with 100 μM NiSO_4_ in 5% BSA/PBS. After another washing step, protein dilutions were added to achieve 600 molecules/µm^2^ ICAM1 and 100 molecules/µm^2^ PfRh5, with or without 100 molecules/µm^2^ of the indicated RIFIN as determined by flow cytometry with bead-supported bilayers. The protein mixtures were incubated for 20 minutes to allow for attachment and then washed. Human anti-RH5 antibody was added at 2 μg/ml for 20 minutes followed by washing. Then, 10^6^ NK cells, isolated from fresh blood samples using a RosetteSep Human NK Cell Enrichment Cocktail (StemCell Technologies), were infused into each lane, followed by incubation for 30 minutes at 37°C. Bilayers were then fixed for 5 minutes in 4% PFA/HBSS followed by washing. Finally, perforin staining was performed using monoclonal anti-perforin-AlexFluor488 (clone dG9, BioLegend) at a concentration of 10 μg/ml for at least an hour. Bilayers were washed three times prior to image acquisition.

Imaging was performed as described before^3^, using an Olympus cell TIRF-4Line system with a 150x (NA 1.45) oil objective at room temperature. Image analysis was performed using ImageJ (v1.54b, NIH) and cell boundaries defined based on segmented (‘default’ algorithm in ImageJ) interference-reflection images ^47^

### Homology Modelling KIR2DS1

The SWISS-MODEL ^48^ web interface was used to generate a template homology model of KIR2DS1 for use in structural comparisons. The protein sequence coding for domains D1 and D2 of KIR2DS1 were input into the tool with a PDB template supplied of KIR2DL1 from our crystallographic data.

## Acknowledgements

This work was funded by a Wellcome Trust collaborative Award (224343/Z/21/Z), Japan Agency for Medical Research and Development (223fa627002h, 24gm1810006h), Japan Science and Technology (CREST, JPMJCR23B5), and Japan Society for the Promotion of Science (22H04989). In addition, this work was supported by the Nippon Foundation. The authors would like to thank Dr Ed Lowe and the beamlines scientists at ESRF beamline ID30A-3 for help with crystallographic data collection.

## Author Contributions

A.S., S.G.C., H.A, M.K.H., and S.I. designed and conceived the study. A.S., H.A., and S.I. identified RBK21 and conducted the functional analysis. A.S. and S.I created RIFIN-expression library and identified KIR2DL1-binding RIFINs from the library. T.E.H. produced protein. S.G.C., and M.K.H. determined the structure of RBK21-KIR2DL1 complex and S.G.C. conducted all other structural and biophysical analysis. A.M.M and M.L.D performed SLB assay of RBK21. A.S., S.G.C., H.A, M.K.H., and S.I. wrote the manuscript and all authors contributed to its revision.

## Conflicts of interest

The authors have no conflicts of interest to declare.

## Data availability

Data within graphs (source data) and uncropped gel and blot images are included with this submission. Crystallographic data is deposited in the protein data bank with accession code 9F2D. All obtained sequence data related with rif-lib1 and -lib2 were deposited at DNA Data Bank of Japan with accession number DRA018487.

## Figures and Figure legends

**Extended Data Figure 1.**
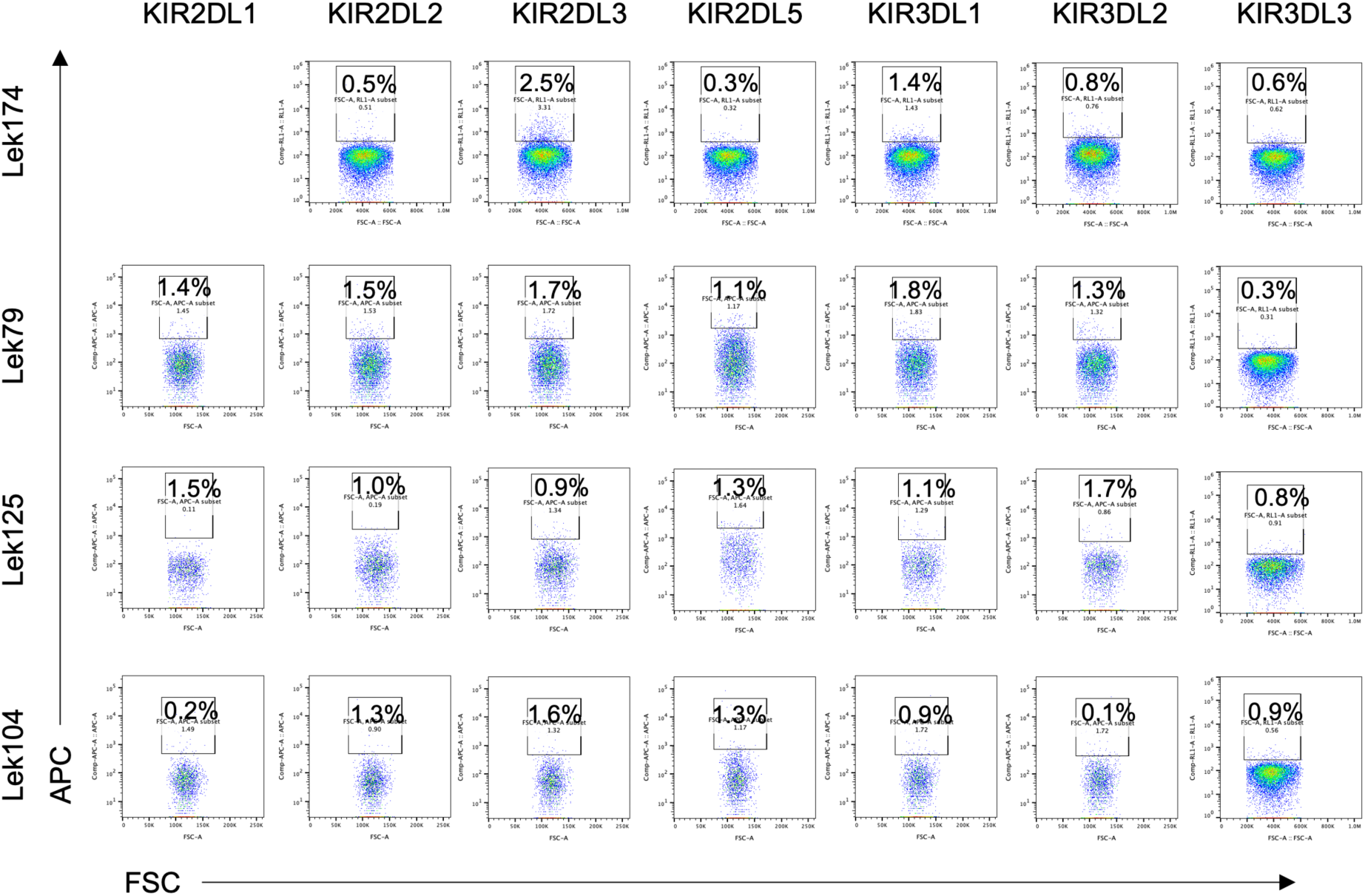
Screening of field-isolated parasites that bind to inhibitory KIR receptors. Percentage of positive iRBCs are shown.

**Extended Data figure 2.**
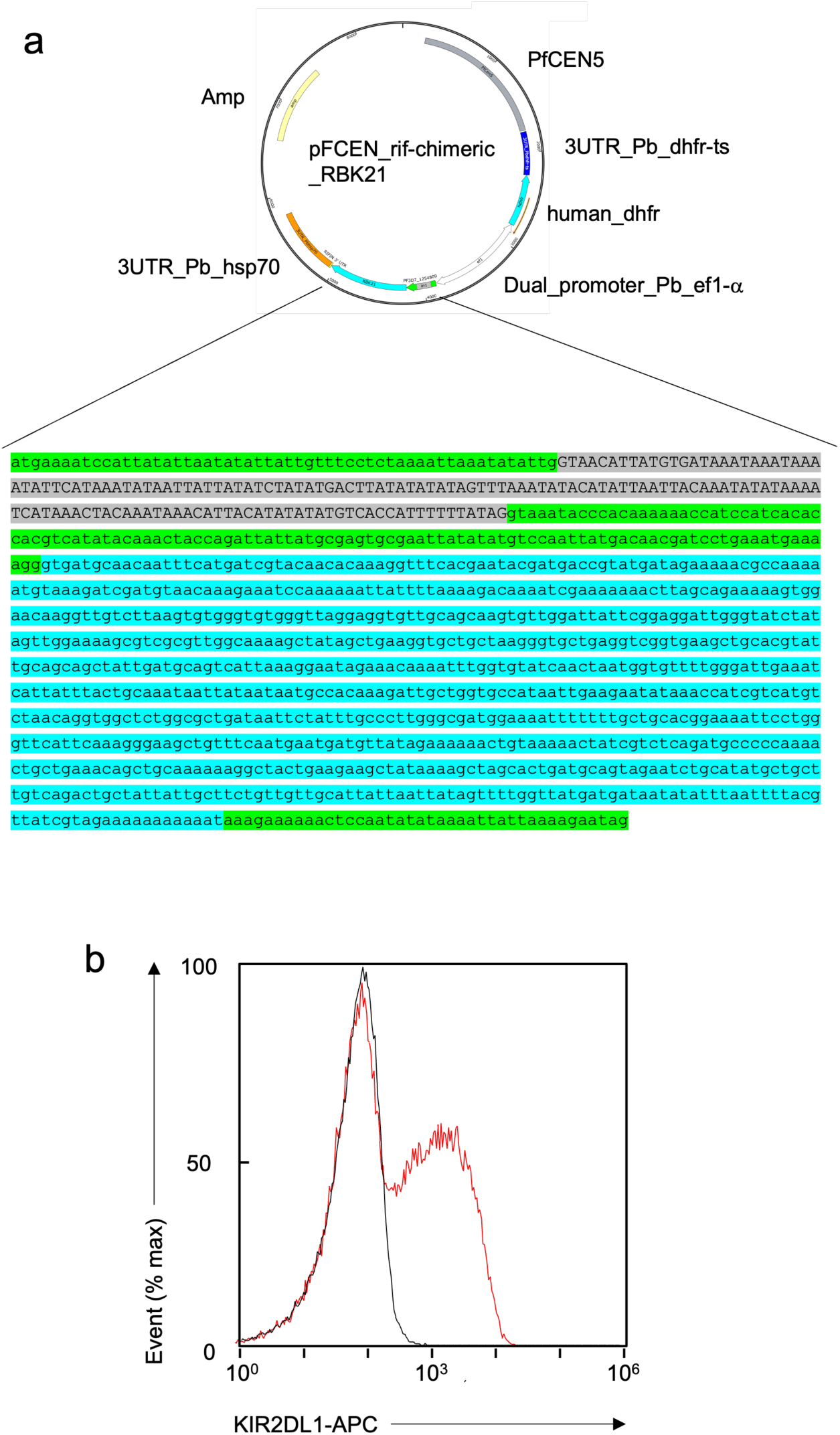
Generation and screening of RIFIN libraries. **a)** Plasmid map of pFCEN-rif featuring the chimeric RBK21 gene. The chimeric RBK21 gene and the human dihydrofolate reductase gene are transcribed under the control of the elongation factor α promoter from *P. berghei.* This promoter has the capability to transcribe genes bidirectionally. In addition, this plasmid has the centromere from chromosome 5 of *P. falciparum* and is maintained stably in the parasite because of function of centromere. The DNA sequence of chimeric RBK21 is shown. The green and blue indicate the conserved region of PF3D7_1254800 and the variable region of RBK21, respectively. Gray shows the intron sequence of PF3D7_1254800 and would be splice out from mature mRNA of chimeric RBK21. **b)** The binding of iRBC with transgenic parasite expressing full sequence of RBK21 was assessed using KIR2DL1-Fc. Black and red show the interaction of KIR2DL1-Fc with *P. falciparum* 3D7 strain and transgenic parasite expressing full RBK21 respectively.

**Extended Data figure 3.**
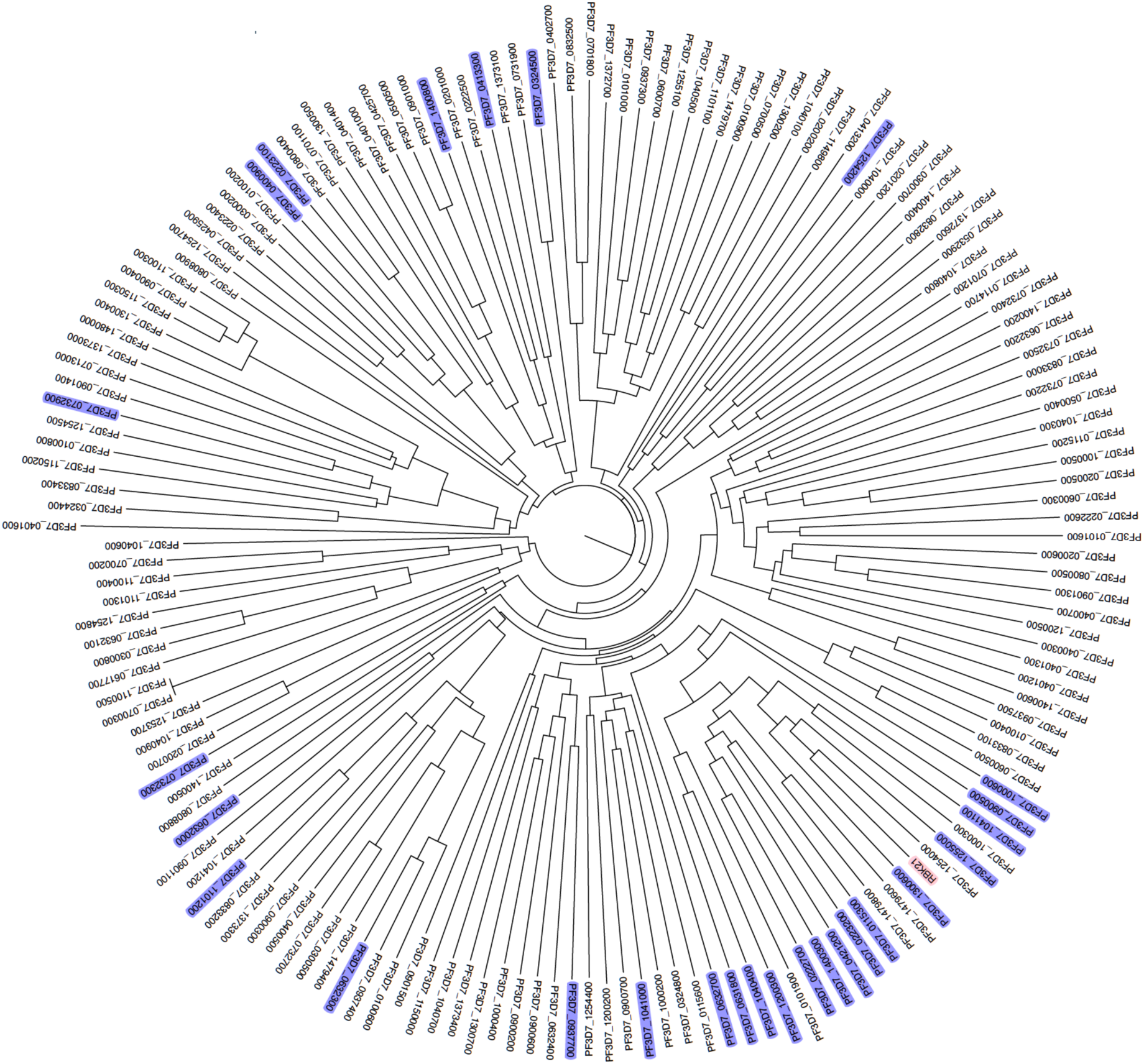
Phylogenetic tree of variable regions of RIFIN of P. falciparum 3D7. Violet and shows KIR2DL1-binding RIFINs while pink highlights RBK21. Those candidates were identified by screening rif-lib1 and -lib2 using KIR2DL1-Fc fusion protein.

**Extended Data figure 4.**
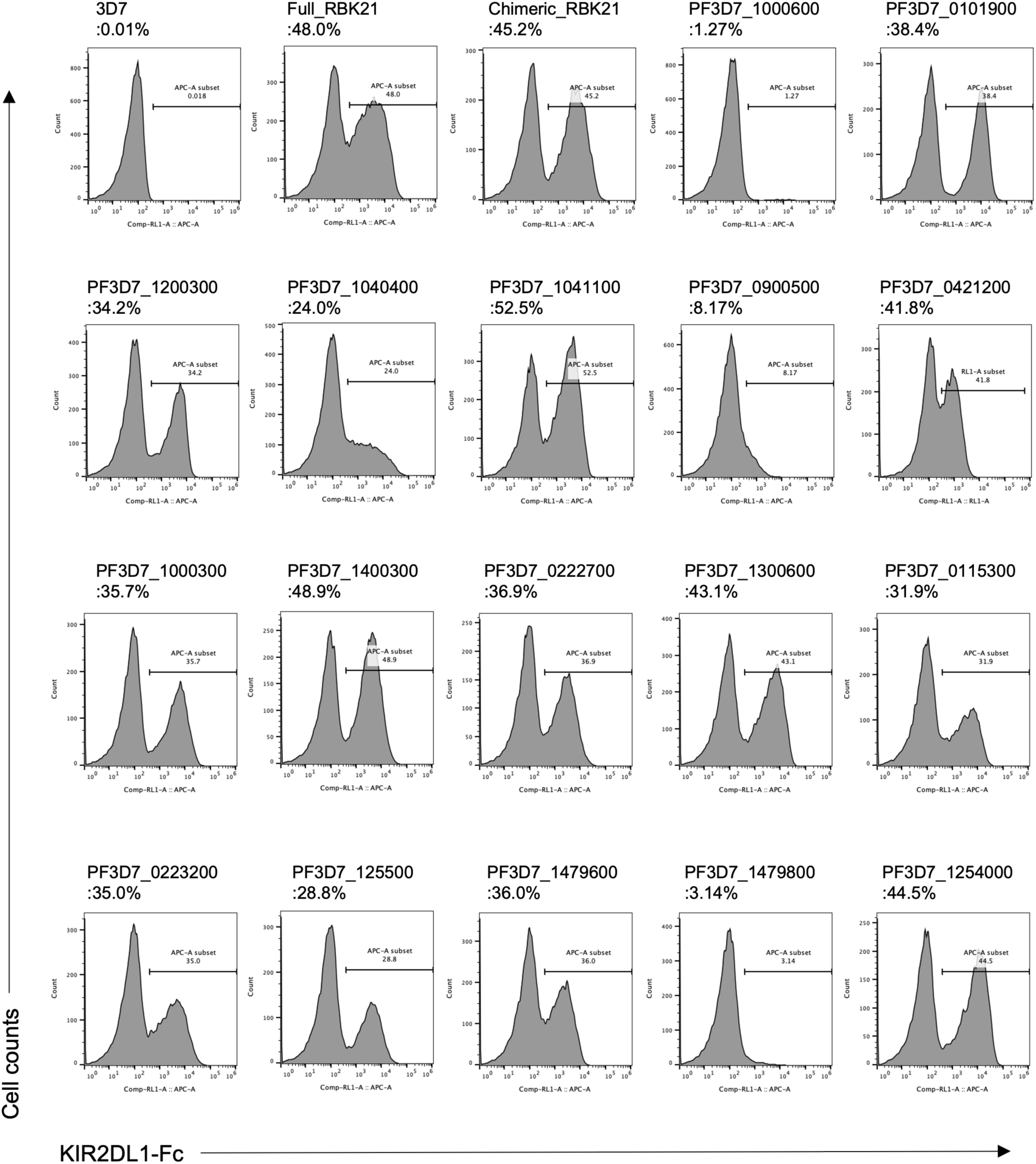
Binding analysis of KIR2DL1-binding RIFINs using the KIR2DL1-Fc fusion protein. All tested RIFINs are classified in the same clade in phylogenetic tree (Fig.1 and Extended data Fig.3). The *P. falciparum* 3D7 strain was used as a negative control. The percentage of positive iRBC is shown at the top of each panel.

**Extended Data Figure 5:**
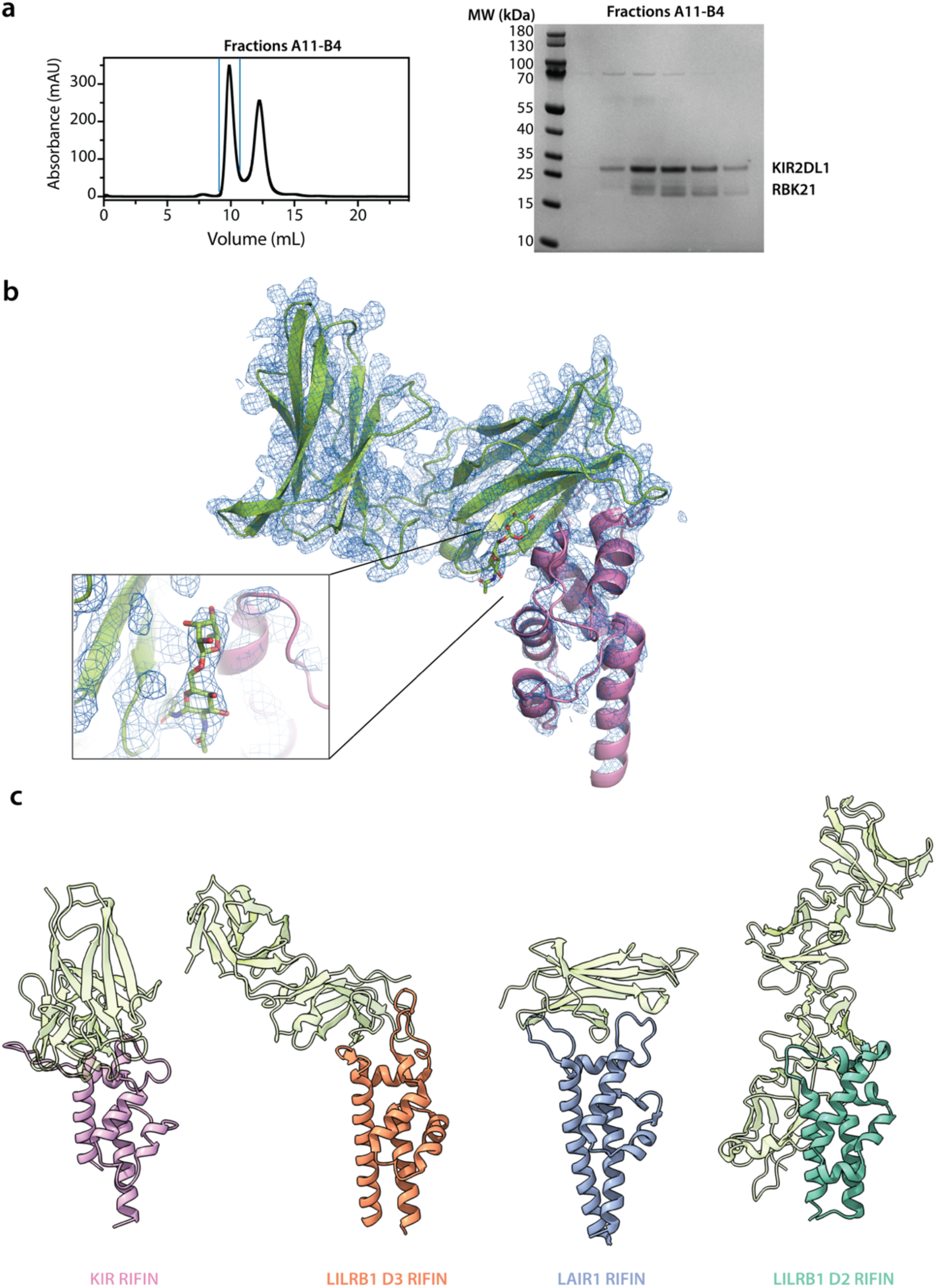
Determination of the structure of RBK21 bound to KIR2DL1 and its comparison with structures of other RIFIN-receptor complexes. **a)** Elution profile from purification of the RBK21-KIR2DL1 complex on a S75 size exclusion chromatography column (left). The fractions corresponding to the peak at 9.8 mL (between the blue lines) were analysed by SDS-PAGE (right). **b)** Overview of the electron density of the complex (blue mesh), with a box, inset, showing the glycan core attached to N67 which is sandwiched at the interface between the RBK21 and the KIR. **c)** A comparison of the structure of solved RIFIN structures, bound to immune receptor ligands shown in light green (PDB codes 7KFK, 7JZK, 6ZDX, respectively), RMSDs between the KIR RIFIN and each other RIFIN across all pairs of backbone atoms were 4.46, 6.68 and 15.37 Å, respectively.

**Extended Data Figure 6:**
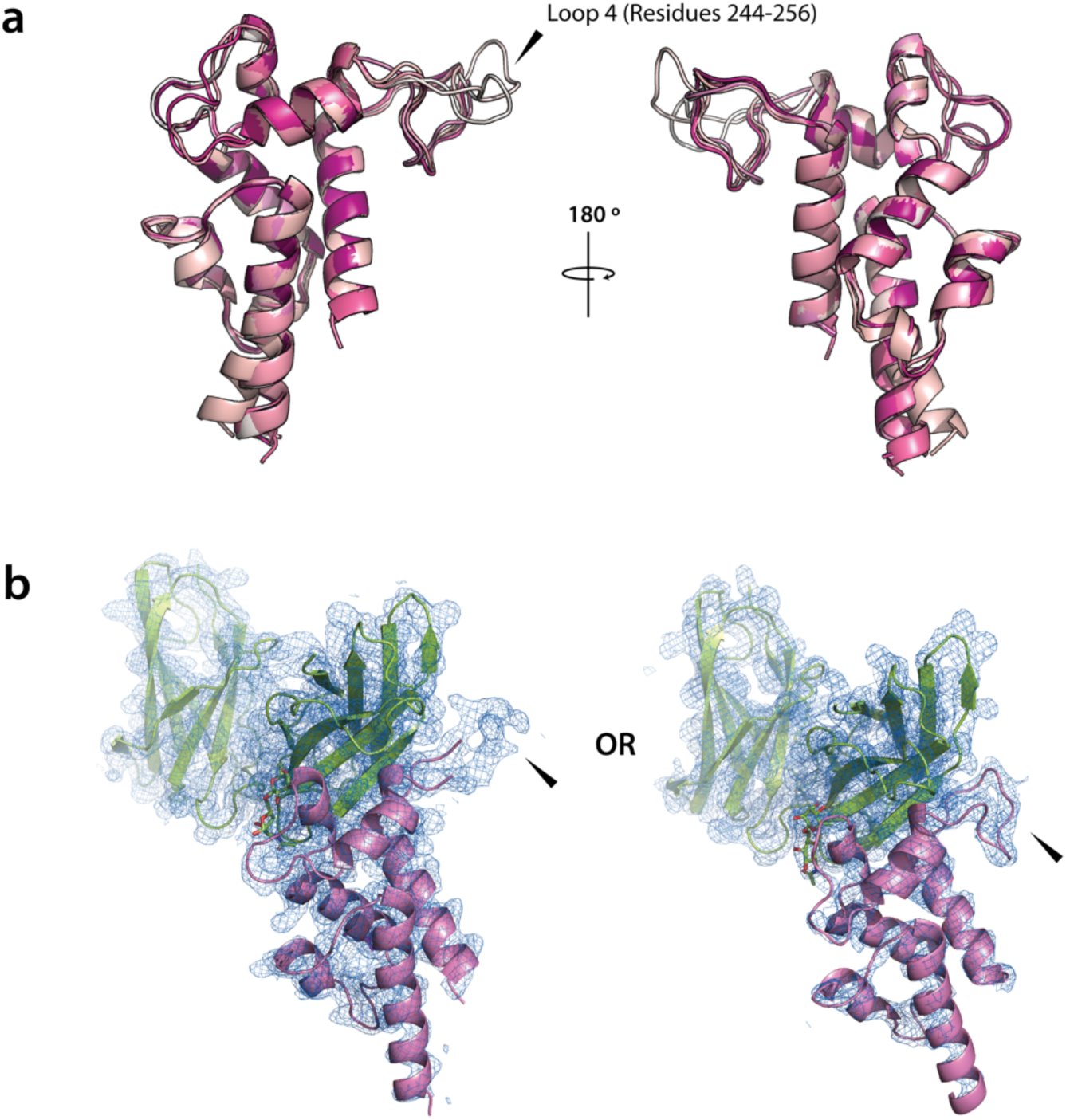
Comparison of RBK21 structures in the crystal asymmetric unit reveal a flexible loop. **a)** Overlay of each copy of RBK21 in the unit cell of the crystal lattice (coloured shades of pink). **b)** Electron density for representatives of the two main conformers of RBK21 with the loop region between residues 244 – 256 in either an upward or downward conformation (black arrow).

**Extended Data Figure 7:**
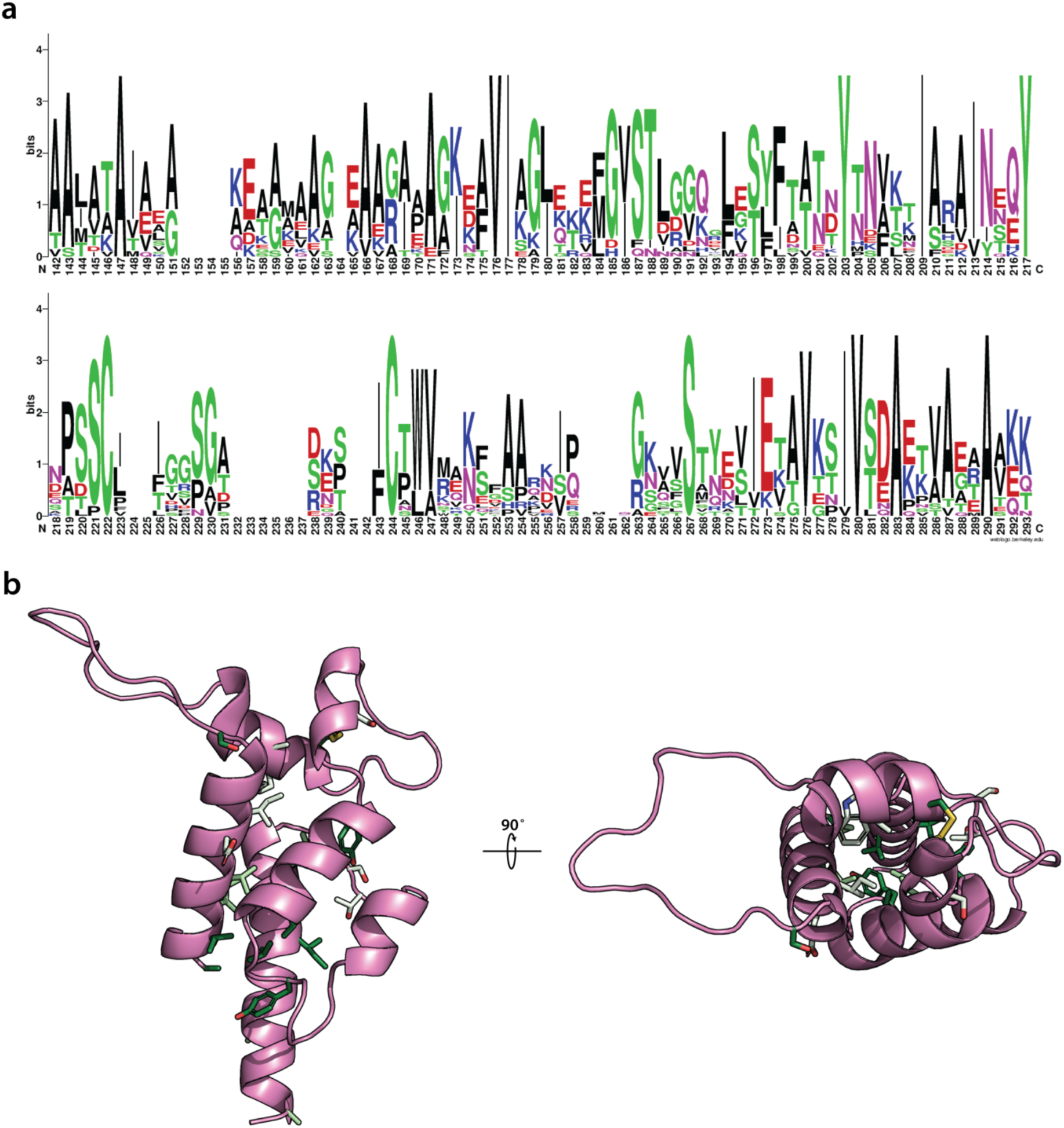
KIR2DL1-binding RIFINs show sequence conservation in their core and diverse surfaces. **a)** Sequence Logo of the identified clade of 3D7 RIFINs that bind to KIR2DL1. Logo was generated using WebLOGO webserver. **b)** Shannon-entropy of sequence conservation was calculated using the Protein Variability Server (PVS) and the most conserved residues mapped onto the protein structure. Entropy values are represented across a gradient of green, with the darkest green representing Entropy values of 0 (absolutely conserved) followed by values between 0 – 0.5 and 0.5 – 0.8. Values above 0.8, representing largescale diversity at the position are not mapped onto the structure.

**Extended Data Figure 8.**
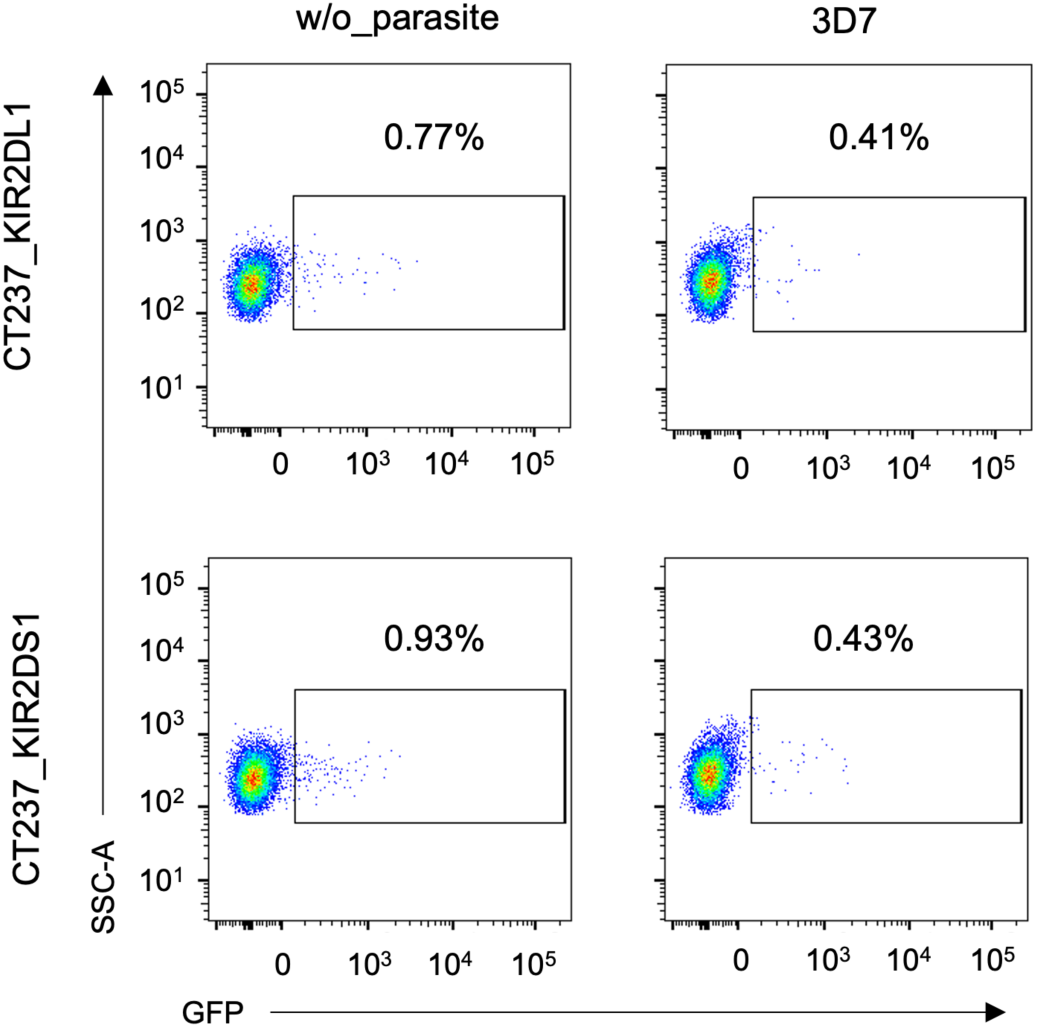
GFP expression in reporter cell lines, which express KIR2DL1 or KIR2DS1 on their surface, was monitored upon stimulation with iRBCs of the *P. falciparum* 3D7 strain. These iRBCs were unable to transduce signals through KIR2DL1 and KIR2DS1.

**Extended Data Figure 9.**
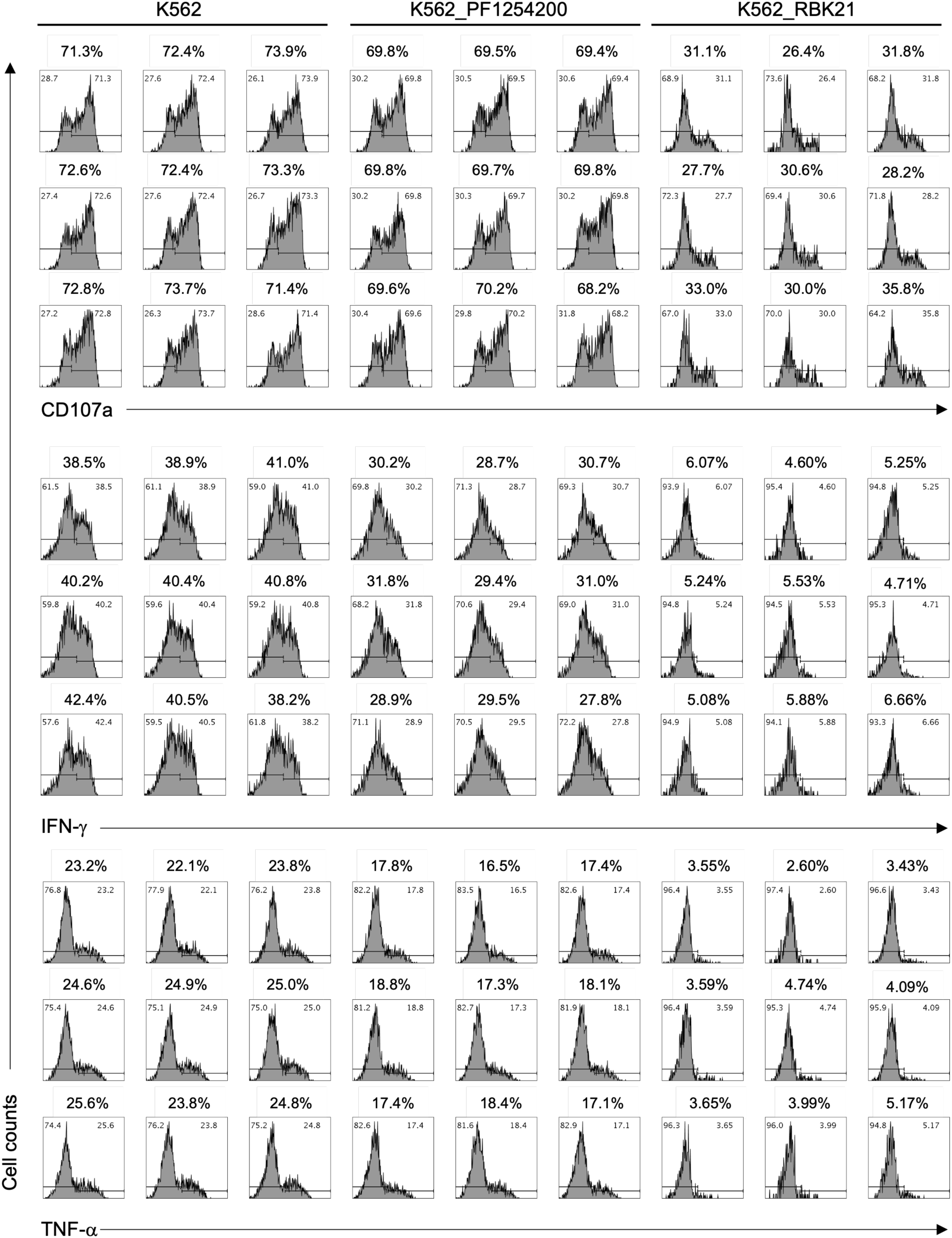
Inhibitory effect of RBK21 on expression CD107a, and production of IFNg and TNFa. Percentage of positive cells in each assay were indicated at the top of panels. K562 and K562 expressing PF3D71254200 (control RIFIN) were used as negative controls. All assays were performed with nine technical replicates.

**Extended Data Figure 10.**
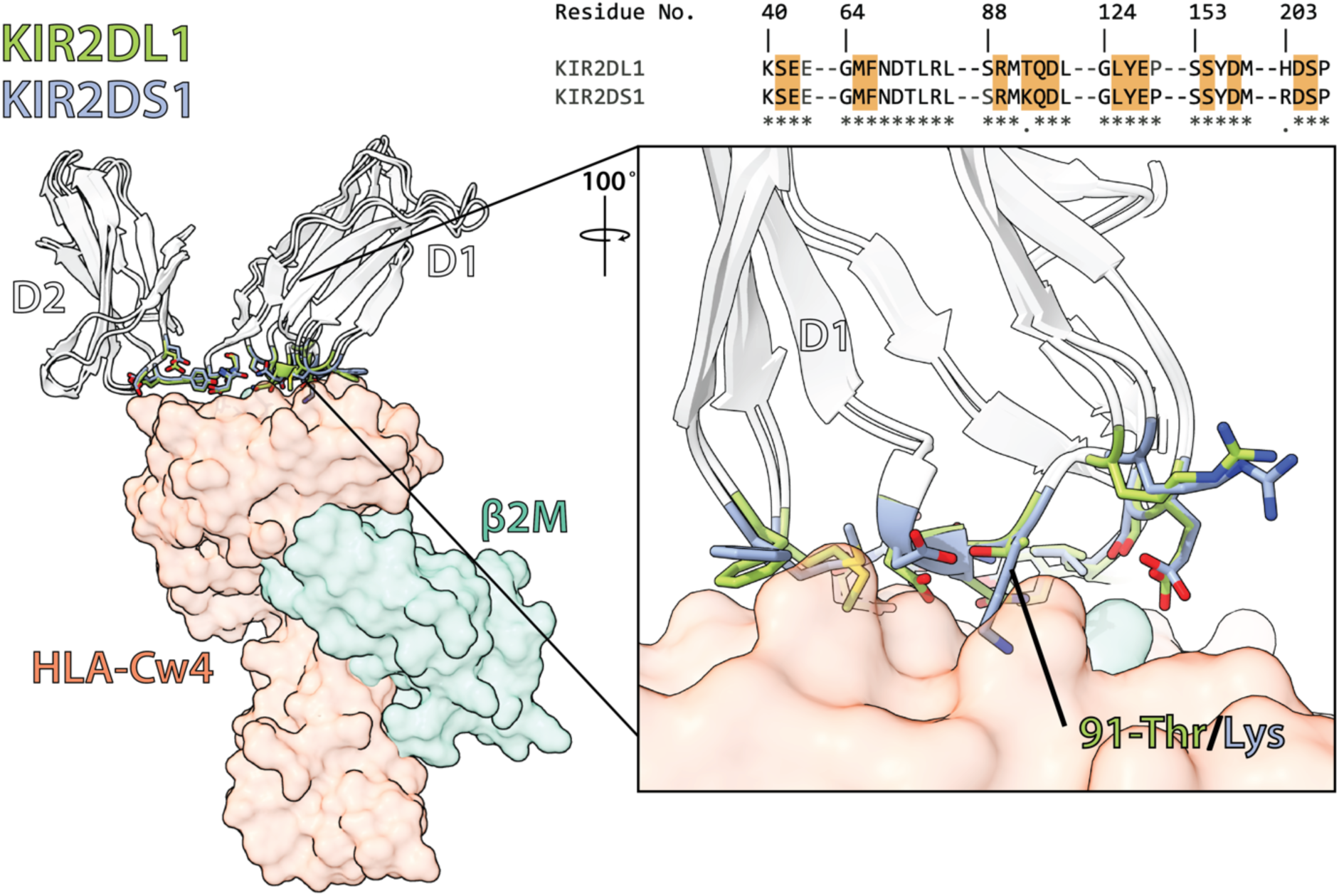
Comparison of the HLA-Cw4 interface between KIR2DL1 and KIR2DS1. **The structure of** KIR2DL1 bound to HLA-Cw4 (PDB 1IM9) was overlaid with a homology model (SWISS-MODEL) of KIR2DS1. Interfacial residues are shown as sticks for KIR2DL1 (green) and KIR2DS2 (blue). The 91Thr/Lys mutation at this interface is labelled. Also shown is a multiple sequence alignment of the regions responsible for the interaction, with residues involved in interactions highlighted in orange (above inset).

**Extended Data Figure 11.**
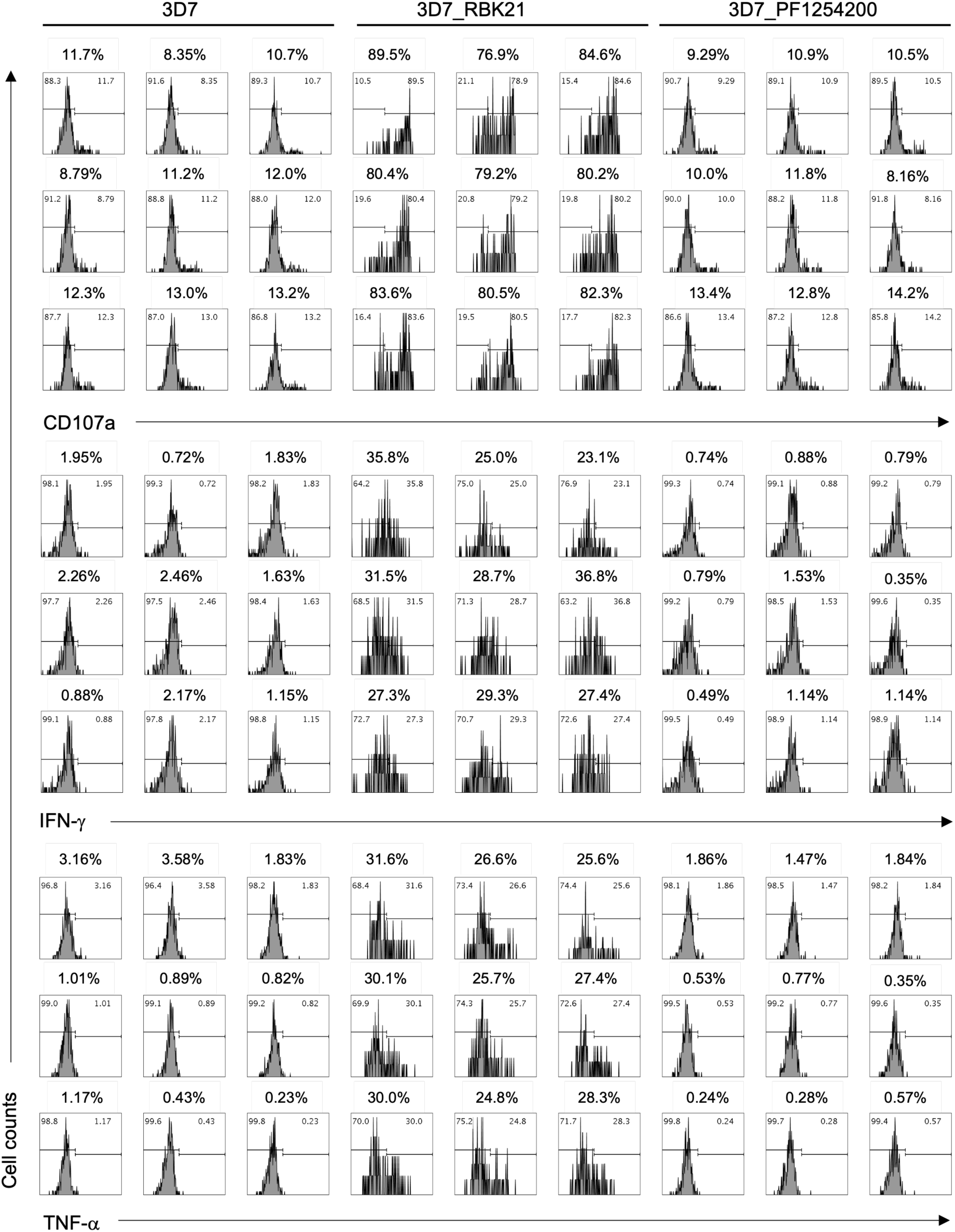
Activation of KIR2DS1-positive NK cells by RBK21. The expression of CD107a, as well as the production of IFN-γ and TNF-α in KIR2DS1-positive NK cells, was monitored. The percentage of positive cells in each assay is indicated at the top of the panels. The iRBCs infected with the *P. falciparum* 3D7 strain and transgenic parasite expressing PF3D71254200 (control RIFIN) were used as negative controls. All assays were performed with nine technical replicates.

**Extended Data Figure 12.**
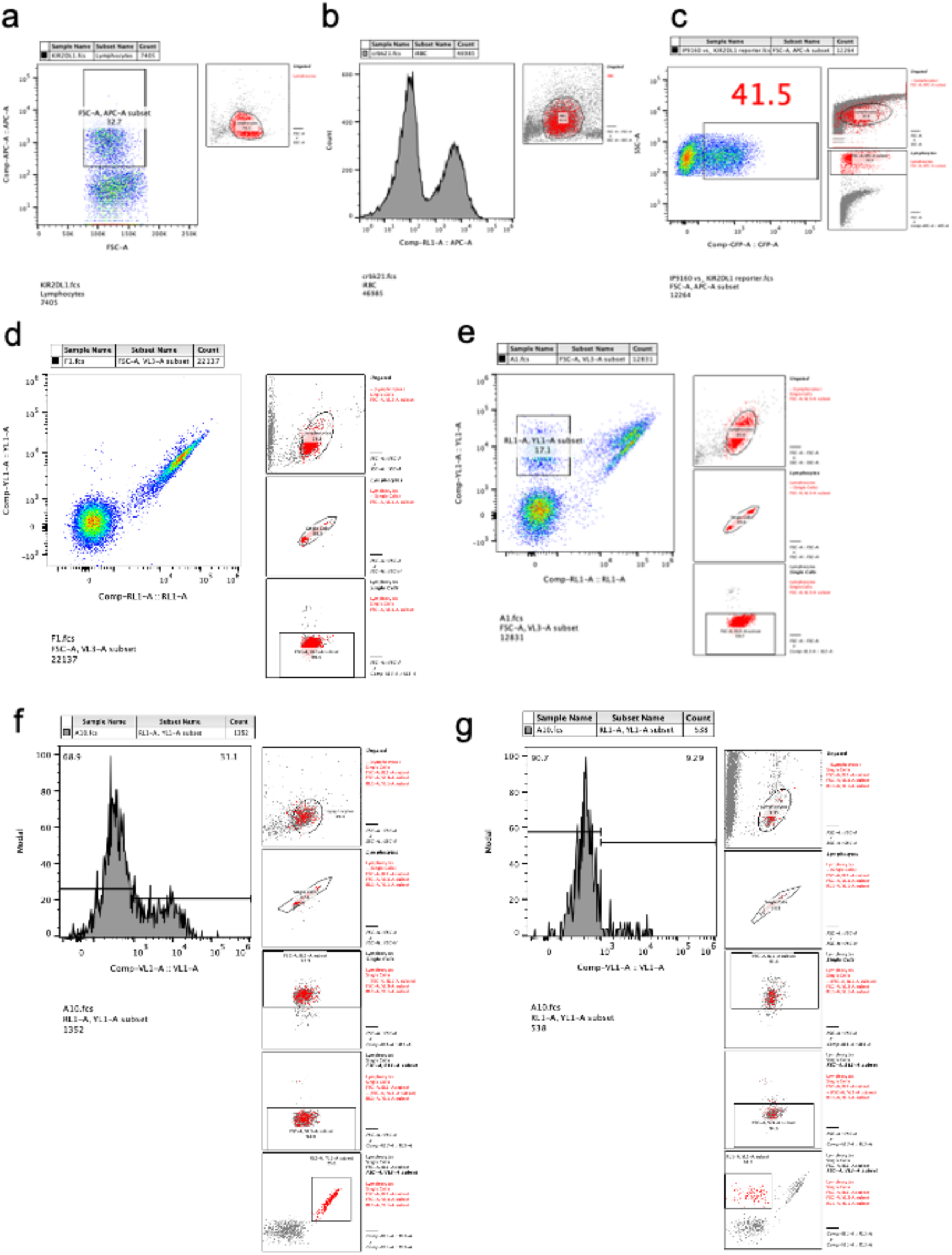
Gating strategies used for this study. **a)** Identification of KIR2DL1-positive iRBCs. This was used for Fig 1a and 1d. b) The binding analysis of transgenic parasites to KIRs-Fc fusion proteins. This was used for Fig. 1b and Extended data Figure 4. c) Detection of GFP positive cell in NFAT-reporter cells. This was used for Figure 3a and 4e. d) Detection of KIR2DL1-positive NK cells in PBMC. This was used for Figure 3b. e) Detection of KIR2DS1-positive NK cells in PBMC. This was used for Figure 4f. f) Detection CD107a positive cells in KIR2DL1-positive NK cells. This was used for Extended data Figure 9. Similar gating strategies were used for detection of cells expressing IFN-g and TNF-a in Extended data Figure 9. g) Detection CD107a positive cells in KIR2DS1-positive NK cells. This was used for Extended data Figure 10. Similar gating strategies were used for detection of IFN-g and TNF-a in the cells in Extended data Figure 10.

**Extended Data Table 1:** Coverage of rif-lib1 for all RIFINs.

| gene_id | annotation | mapped_reads_rif_lib1_1<br>_before | normalization_rif_lib1_1-<br>1_before |
| --- | --- | --- | --- |
| PF3D7_1000600 | rifin | 0 | 0 |
| PF3D7_0401600 | rifin | 0 | 0 |
| PF3D7_0901100 | rifin | 0 | 0 |
| PF3D7_1253700 | rifin | 0 | 0 |
| PF3D7_0832500 | rifin | 0 | 0 |
| PF3D7_0701800 | rifin | 0 | 0 |
| PF3D7_0700200 | rifin | 0 | 0 |
| PF3D7_0632000 | rifin | 1 | 4.97958E-06 |
| PF3D7_1100400 | rifin | 1 | 4.97958E-06 |
| PF3D7_0732900 | rifin | 1 | 4.97958E-06 |
| PF3D7_0617700 | rifin | 2 | 9.95917E-06 |
| PF3D7_1400400 | rifin | 3 | 1.49388E-05 |
| PF3D7_0600500 | rifin | 3 | 1.49388E-05 |
| PF3D7_0600700 | rifin | 3 | 1.49388E-05 |
| PF3D7_0100400 | rifin | 3 | 1.49388E-05 |
| PF3D7_1300700 | rifin | 3 | 1.49388E-05 |
| PF3D7_1150300 | rifin | 4 | 1.99183E-05 |
| PF3D7_0425900 | rifin | 4 | 1.99183E-05 |
| PF3D7_0413200 | rifin | 5 | 2.48979E-05 |
| PF3D7_1150200 | rifin | 5 | 2.48979E-05 |
| PF3D7_1000400 | rifin | 5 | 2.48979E-05 |
| PF3D7_1254400 | rifin | 6 | 2.98775E-05 |
| PF3D7_0300700 | rifin | 6 | 2.98775E-05 |
| PF3D7_0223100 | rifin | 7 | 3.48571E-05 |
| PF3D7_1479600 | rifin | 7 | 3.48571E-05 |
| PF3D7_0833000 | rifin | 7 | 3.48571E-05 |
| PF3D7_0808800 | rifin | 8 | 3.98367E-05 |
| PF3D7_0100900 | rifin | 8 | 3.98367E-05 |
| PF3D7_0713000 | rifin | 9 | 4.48163E-05 |
| PF3D7_1479700 | rifin | 9 | 4.48163E-05 |
| PF3D7_0100600 | rifin | 10 | 4.97958E-05 |
| PF3D7_0732500 | rifin | 11 | 5.47754E-05 |
| PF3D7_0833200 | rifin | 11 | 5.47754E-05 |
| PF3D7_1040500 | rifin | 12 | 5.9755E-05 |
| PF3D7_0200700 | rifin | 12 | 5.9755E-05 |
| PF3D7_0324800 | rifin | 14 | 6.97142E-05 |
| PF3D7_0800400 | rifin | 14 | 6.97142E-05 |
| PF3D7_0732400 | rifin | 15 | 7.46938E-05 |
| PF3D7_0900200 | rifin | 15 | 7.46938E-05 |
| PF3D7_0732700 | rifin | 17 | 8.46529E-05 |
| PF3D7_0400900 | rifin | 17 | 8.46529E-05 |
| PF3D7_0300200 | rifin | 18 | 8.96325E-05 |
| PF3D7_0937700 | rifin | 20 | 9.95917E-05 |
| PF3D7_0901500 | rifin | 20 | 9.95917E-05 |
| PF3D7_1101100 | rifin | 23 | 0.00011453 |
| PF3D7_0937400 | rifin | 23 | 0.00011453 |
| PF3D7_0400700 | rifin | 23 | 0.00011453 |
| PF3D7_0223400 | rifin | 24 | 0.00011951 |
| PF3D7_1040800 | rifin | 27 | 0.000134449 |
| PF3D7_0200600 | rifin | 29 | 0.000144408 |
| PF3D7_1200500 | rifin | 29 | 0.000144408 |
| PF3D7_1040000 | rifin | 30 | 0.000149388 |
| PF3D7_1479400 | rifin | 30 | 0.000149388 |
| PF3D7_0900500 | rifin | 31 | 0.000154367 |
| PF3D7_1254700 | rifin | 31 | 0.000154367 |
| PF3D7_0900300 | rifin | 31 | 0.000154367 |
| PF3D7_1400200 | rifin | 36 | 0.000179265 |
| PF3D7_1101300 | rifin | 37 | 0.000184245 |
| PF3D7_0300800 | rifin | 37 | 0.000184245 |
| PF3D7_1041200 | rifin | 37 | 0.000184245 |
| PF3D7_0632400 | rifin | 38 | 0.000189224 |
| PF3D7_0114700 | rifin | 43 | 0.000214122 |
| PF3D7_1300200 | rifin | 45 | 0.000224081 |
| PF3D7_0632300 | rifin | 45 | 0.000224081 |
| PF3D7_0833100 | rifin | 45 | 0.000224081 |
| PF3D7_0401200 | rifin | 46 | 0.000229061 |
| PF3D7_1100300 | rifin | 53 | 0.000263918 |
| PF3D7_1300400 | rifin | 54 | 0.000268898 |
| PF3D7_1400800 | rifin | 55 | 0.000273877 |
| PF3D7_0937500 | rifin | 56 | 0.000278857 |
| PF3D7_0400300 | rifin | 58 | 0.000288816 |
| PF3D7_1255100 | rifin | 59 | 0.000293795 |
| PF3D7_1041000 | rifin | 60 | 0.000298775 |
| PF3D7_0632200 | rifin | 60 | 0.000298775 |
| PF3D7_0632100 | rifin | 62 | 0.000308734 |
| PF3D7_0900700 | rifin | 64 | 0.000318693 |
| PF3D7_1000500 | rifin | 68 | 0.000338612 |
| PF3D7_1480000 | rifin | 70 | 0.000348571 |
| PF3D7_0901300 | rifin | 71 | 0.00035355 |
| PF3D7_0400500 | rifin | 75 | 0.000373469 |
| PF3D7_1400500 | rifin | 76 | 0.000378448 |
| PF3D7_0300500 | rifin | 76 | 0.000378448 |
| PF3D7_0425700 | rifin | 78 | 0.000388408 |
| PF3D7_0900400 | rifin | 81 | 0.000403346 |
| PF3D7_1000200 | rifin | 82 | 0.000408326 |
| PF3D7_0222600 | rifin | 83 | 0.000413305 |
| PF3D7_1101200 | rifin | 84 | 0.000418285 |
| PF3D7_1373100 | rifin | 86 | 0.000428244 |
| PF3D7_0800500 | rifin | 97 | 0.00048302 |
| PF3D7_1254200 | rifin | 111 | 0.000552734 |
| PF3D7_0222700 | rifin | 118 | 0.000587591 |
| PF3D7_0413300 | rifin | 120 | 0.00059755 |
| PF3D7_0632700 | rifin | 127 | 0.000632407 |
| PF3D7_0701100 | rifin | 127 | 0.000632407 |
| PF3D7_0200500 | rifin | 129 | 0.000642366 |
| PF3D7_0101900 | rifin | 129 | 0.000642366 |
| PF3D7_0101600 | rifin | 161 | 0.000801713 |
| PF3D7_1372700 | rifin | 162 | 0.000806693 |
| PF3D7_1372600 | rifin | 163 | 0.000811672 |
| PF3D7_1254800 | rifin | 183 | 0.000911264 |
| PF3D7_0808900 | rifin | 188 | 0.000936162 |
| PF3D7_0115600 | rifin | 198 | 0.000985958 |
| PF3D7_0732200 | rifin | 218 | 0.001085549 |
| PF3D7_0201200 | rifin | 221 | 0.001100488 |
| PF3D7_0937300 | rifin | 261 | 0.001299671 |
| PF3D7_0100200 | rifin | 279 | 0.001389304 |
| PF3D7_0115300 | rifin | 282 | 0.001404243 |
| PF3D7_0700500 | rifin | 292 | 0.001454038 |
| PF3D7_0100800 | rifin | 318 | 0.001583508 |
| PF3D7_0901400 | rifin | 349 | 0.001737875 |
| PF3D7_0200200 | rifin | 376 | 0.001872323 |
| PF3D7_1200200 | rifin | 411 | 0.002046609 |
| PF3D7_0222500 | rifin | 419 | 0.002086446 |
| PF3D7_0401300 | rifin | 460 | 0.002290609 |
| PF3D7_1040700 | rifin | 497 | 0.002474853 |
| PF3D7_1040400 | rifin | 501 | 0.002494771 |
| PF3D7_0500400 | rifin | 531 | 0.002644159 |
| PF3D7_1040300 | rifin | 535 | 0.002664077 |
| PF3D7_1300600 | rifin | 544 | 0.002708894 |
| PF3D7_1255000 | rifin | 587 | 0.002923016 |
| PF3D7_0600300 | rifin | 596 | 0.002967832 |
| PF3D7_1400600 | rifin | 670 | 0.003336321 |
| PF3D7_0201000 | rifin | 712 | 0.003545464 |
| PF3D7_0115200 | rifin | 783 | 0.003899014 |
| PF3D7_0732300 | rifin | 864 | 0.00430236 |
| PF3D7_0401400 | rifin | 960 | 0.0047804 |
| PF3D7_0421200 | rifin | 994 | 0.004949706 |
| PF3D7_1041100 | rifin | 999 | 0.004974604 |
| PF3D7_0324500 | rifin | 1220 | 0.006075092 |
| PF3D7_1373300 | rifin | 1239 | 0.006169704 |
| PF3D7_1100500 | rifin | 1296 | 0.00645354 |
| PF3D7_1149800 | rifin | 1296 | 0.00645354 |
| PF3D7_1300500 | rifin | 1429 | 0.007115825 |
| PF3D7_1373000 | rifin | 1590 | 0.007917538 |
| PF3D7_0101000 | rifin | 1652 | 0.008226272 |
| PF3D7_0631800 | rifin | 1687 | 0.008400558 |
| PF3D7_1400300 | rifin | 2228 | 0.011094512 |
| PF3D7_0900600 | rifin | 2303 | 0.011467981 |
| PF3D7_0833400 | rifin | 2682 | 0.013355244 |
| PF3D7_0223200 | rifin | 2746 | 0.013673937 |
| PF3D7_1040900 | rifin | 2768 | 0.013783488 |
| PF3D7_1479800 | rifin | 3151 | 0.015690668 |
| PF3D7_1200300 | rifin | 3171 | 0.01579026 |
| PF3D7_0324400 | rifin | 4012 | 0.01997809 |
| PF3D7_0700300 | rifin | 4129 | 0.020560701 |
| PF3D7_0401000 | rifin | 4329 | 0.021556618 |
| PF3D7_1150000 | rifin | 4504 | 0.022428045 |
| PF3D7_1373400 | rifin | 5592 | 0.027845832 |
| PF3D7_0901000 | rifin | 6075 | 0.030250971 |
| PF3D7_1040600 | rifin | 6133 | 0.030539787 |
| PF3D7_1254000 | rifin | 7999 | 0.03983169 |
| PF3D7_1000300 | rifin | 8585 | 0.042749726 |
| PF3D7_1254500 | rifin | 10859 | 0.054073299 |
| PF3D7_0532900 | rifin | 15032 | 0.074853102 |
| PF3D7_0832800 | rifin | 16214 | 0.08073897 |
| PF3D7_1040100 | rifin | 20181 | 0.100492979 |
| PF3D7_0500500 | rifin | 24641 | 0.122701922 |

**Extended Data Table 2:** Coverage of rif-lib2 for all RIFINs.

| gene_id | annotation | mapped_reads_rif_lib1_2<br>_before | normalization_rif_lib_1-<br>2_before |
| --- | --- | --- | --- |
| PF3D7_0401600 | rifin | 0 | 0 |
| PF3D7_1253700 | rifin | 0 | 0 |
| PF3D7_0832500 | rifin | 0 | 0 |
| PF3D7_0700200 | rifin | 0 | 0 |
| PF3D7_1000600 | rifin | 4 | 4.74126E-06 |
| PF3D7_0901100 | rifin | 4 | 4.74126E-06 |
| PF3D7_0632000 | rifin | 5 | 5.92658E-06 |
| PF3D7_1400400 | rifin | 5 | 5.92658E-06 |
| PF3D7_0732900 | rifin | 5 | 5.92658E-06 |
| PF3D7_0617700 | rifin | 8 | 9.48253E-06 |
| PF3D7_1254400 | rifin | 9 | 1.06678E-05 |
| PF3D7_1479600 | rifin | 9 | 1.06678E-05 |
| PF3D7_0425900 | rifin | 11 | 1.30385E-05 |
| PF3D7_1150300 | rifin | 12 | 1.42238E-05 |
| PF3D7_0808800 | rifin | 13 | 1.54091E-05 |
| PF3D7_1000400 | rifin | 17 | 2.01504E-05 |
| PF3D7_1300700 | rifin | 17 | 2.01504E-05 |
| PF3D7_0413200 | rifin | 21 | 2.48916E-05 |
| PF3D7_0701800 | rifin | 21 | 2.48916E-05 |
| PF3D7_0200700 | rifin | 22 | 2.60769E-05 |
| PF3D7_0300200 | rifin | 30 | 3.55595E-05 |
| PF3D7_0223100 | rifin | 31 | 3.67448E-05 |
| PF3D7_0100400 | rifin | 33 | 3.91154E-05 |
| PF3D7_0901500 | rifin | 34 | 4.03007E-05 |
| PF3D7_0200600 | rifin | 34 | 4.03007E-05 |
| PF3D7_1150200 | rifin | 35 | 4.14861E-05 |
| PF3D7_0600500 | rifin | 38 | 4.5042E-05 |
| PF3D7_0800400 | rifin | 40 | 4.74126E-05 |
| PF3D7_0400900 | rifin | 44 | 5.21539E-05 |
| PF3D7_0937700 | rifin | 44 | 5.21539E-05 |
| PF3D7_0100900 | rifin | 44 | 5.21539E-05 |
| PF3D7_0300700 | rifin | 49 | 5.80805E-05 |
| PF3D7_1100400 | rifin | 50 | 5.92658E-05 |
| PF3D7_1479700 | rifin | 61 | 7.23043E-05 |
| PF3D7_0833000 | rifin | 62 | 7.34896E-05 |
| PF3D7_1040500 | rifin | 64 | 7.58602E-05 |
| PF3D7_0324800 | rifin | 66 | 7.82308E-05 |
| PF3D7_0100600 | rifin | 71 | 8.41574E-05 |
| PF3D7_1300200 | rifin | 73 | 8.65281E-05 |
| PF3D7_0900700 | rifin | 76 | 9.0084E-05 |
| PF3D7_0400700 | rifin | 77 | 9.12693E-05 |
| PF3D7_0833200 | rifin | 78 | 9.24546E-05 |
| PF3D7_1040800 | rifin | 79 | 9.364E-05 |
| PF3D7_0223400 | rifin | 81 | 9.60106E-05 |
| PF3D7_1372700 | rifin | 86 | 0.000101937 |
| PF3D7_1300400 | rifin | 95 | 0.000112605 |
| PF3D7_0425700 | rifin | 104 | 0.000123273 |
| PF3D7_1040000 | rifin | 104 | 0.000123273 |
| PF3D7_0732500 | rifin | 105 | 0.000124458 |
| PF3D7_1041000 | rifin | 107 | 0.000126829 |
| PF3D7_0800500 | rifin | 117 | 0.000138682 |
| PF3D7_1000500 | rifin | 120 | 0.000142238 |
| PF3D7_1255100 | rifin | 128 | 0.00015172 |
| PF3D7_1100300 | rifin | 134 | 0.000158832 |
| PF3D7_0937400 | rifin | 135 | 0.000160018 |
| PF3D7_0732400 | rifin | 136 | 0.000161203 |
| PF3D7_1200500 | rifin | 138 | 0.000163574 |
| PF3D7_0833100 | rifin | 139 | 0.000164759 |
| PF3D7_0300800 | rifin | 148 | 0.000175427 |
| PF3D7_0222600 | rifin | 155 | 0.000183724 |
| PF3D7_1480000 | rifin | 157 | 0.000186095 |
| PF3D7_1101300 | rifin | 159 | 0.000188465 |
| PF3D7_0401300 | rifin | 159 | 0.000188465 |
| PF3D7_0400300 | rifin | 169 | 0.000200318 |
| PF3D7_0632400 | rifin | 175 | 0.00020743 |
| PF3D7_1254700 | rifin | 176 | 0.000208616 |
| PF3D7_1101100 | rifin | 178 | 0.000210986 |
| PF3D7_0200500 | rifin | 180 | 0.000213357 |
| PF3D7_1400800 | rifin | 181 | 0.000214542 |
| PF3D7_0632700 | rifin | 190 | 0.00022521 |
| PF3D7_1254200 | rifin | 190 | 0.00022521 |
| PF3D7_0713000 | rifin | 205 | 0.00024299 |
| PF3D7_0900300 | rifin | 218 | 0.000258399 |
| PF3D7_0632300 | rifin | 226 | 0.000267881 |
| PF3D7_0632100 | rifin | 229 | 0.000271437 |
| PF3D7_0900200 | rifin | 242 | 0.000286846 |
| PF3D7_0101600 | rifin | 244 | 0.000289217 |
| PF3D7_0114700 | rifin | 248 | 0.000293958 |
| PF3D7_0222700 | rifin | 249 | 0.000295144 |
| PF3D7_0900500 | rifin | 255 | 0.000302256 |
| PF3D7_1372600 | rifin | 256 | 0.000303441 |
| PF3D7_0732700 | rifin | 257 | 0.000304626 |
| PF3D7_0101900 | rifin | 292 | 0.000346112 |
| PF3D7_0808900 | rifin | 323 | 0.000382857 |
| PF3D7_0900400 | rifin | 336 | 0.000398266 |
| PF3D7_0600700 | rifin | 349 | 0.000413675 |
| PF3D7_0401200 | rifin | 364 | 0.000431455 |
| PF3D7_1041200 | rifin | 372 | 0.000440937 |
| PF3D7_1400500 | rifin | 388 | 0.000459903 |
| PF3D7_0937500 | rifin | 433 | 0.000513242 |
| PF3D7_1000200 | rifin | 465 | 0.000551172 |
| PF3D7_0400500 | rifin | 466 | 0.000552357 |
| PF3D7_0632200 | rifin | 474 | 0.00056184 |
| PF3D7_0413300 | rifin | 485 | 0.000574878 |
| PF3D7_0421200 | rifin | 518 | 0.000613994 |
| PF3D7_1101200 | rifin | 529 | 0.000627032 |
| PF3D7_1254800 | rifin | 571 | 0.000676815 |
| PF3D7_0115600 | rifin | 582 | 0.000689854 |
| PF3D7_0201200 | rifin | 634 | 0.00075149 |
| PF3D7_1200200 | rifin | 635 | 0.000752676 |
| PF3D7_0701100 | rifin | 742 | 0.000879504 |
| PF3D7_0901300 | rifin | 775 | 0.00091862 |
| PF3D7_0937300 | rifin | 878 | 0.001040707 |
| PF3D7_0300500 | rifin | 886 | 0.00105019 |
| PF3D7_1400200 | rifin | 894 | 0.001059672 |
| PF3D7_0200200 | rifin | 898 | 0.001064414 |
| PF3D7_1479400 | rifin | 931 | 0.001103529 |
| PF3D7_0732200 | rifin | 970 | 0.001149756 |
| PF3D7_0700500 | rifin | 1039 | 0.001231543 |
| PF3D7_0115300 | rifin | 1096 | 0.001299106 |
| PF3D7_0631800 | rifin | 1104 | 0.001308589 |
| PF3D7_0100800 | rifin | 1284 | 0.001521946 |
| PF3D7_1040700 | rifin | 1289 | 0.001527872 |
| PF3D7_0201000 | rifin | 1416 | 0.001678407 |
| PF3D7_0901400 | rifin | 1485 | 0.001760194 |
| PF3D7_0100200 | rifin | 1563 | 0.001852649 |
| PF3D7_0600300 | rifin | 1638 | 0.001941547 |
| PF3D7_1300500 | rifin | 1707 | 0.002023334 |
| PF3D7_0401400 | rifin | 1829 | 0.002167943 |
| PF3D7_1041100 | rifin | 1968 | 0.002332702 |
| PF3D7_1040300 | rifin | 2185 | 0.002589915 |
| PF3D7_0222500 | rifin | 2717 | 0.003220503 |
| PF3D7_1255000 | rifin | 2795 | 0.003312958 |
| PF3D7_1300600 | rifin | 2912 | 0.00345164 |
| PF3D7_1400600 | rifin | 2996 | 0.003551206 |
| PF3D7_1373000 | rifin | 3256 | 0.003859388 |
| PF3D7_0324500 | rifin | 4075 | 0.004830162 |
| PF3D7_1149800 | rifin | 4249 | 0.005036407 |
| PF3D7_0500400 | rifin | 4331 | 0.005133603 |
| PF3D7_0732300 | rifin | 4498 | 0.005331551 |
| PF3D7_0101000 | rifin | 5022 | 0.005952656 |
| PF3D7_1100500 | rifin | 5038 | 0.005971621 |
| PF3D7_1040900 | rifin | 5403 | 0.006404261 |
| PF3D7_1479800 | rifin | 6228 | 0.007382147 |
| PF3D7_0833400 | rifin | 6771 | 0.008025774 |
| PF3D7_1373100 | rifin | 9219 | 0.010927427 |
| PF3D7_0115200 | rifin | 11508 | 0.013640615 |
| PF3D7_0223200 | rifin | 11566 | 0.013709363 |
| PF3D7_0700300 | rifin | 12621 | 0.014959871 |
| PF3D7_0532900 | rifin | 13185 | 0.015628389 |
| PF3D7_0401000 | rifin | 13672 | 0.016205638 |
| PF3D7_1400300 | rifin | 14685 | 0.017406363 |
| PF3D7_1040400 | rifin | 16001 | 0.018966239 |
| PF3D7_0324400 | rifin | 16002 | 0.018967424 |
| PF3D7_1150000 | rifin | 16359 | 0.019390582 |
| PF3D7_1254000 | rifin | 17444 | 0.020676649 |
| PF3D7_1373300 | rifin | 18037 | 0.021379542 |
| PF3D7_1373400 | rifin | 18502 | 0.021930714 |
| PF3D7_1040600 | rifin | 28582 | 0.033878697 |
| PF3D7_1000300 | rifin | 35883 | 0.042532688 |
| PF3D7_1200300 | rifin | 40562 | 0.048078781 |
| PF3D7_1040100 | rifin | 46124 | 0.054671507 |
| PF3D7_1254500 | rifin | 49887 | 0.059131851 |
| PF3D7_0900600 | rifin | 50923 | 0.060359838 |
| PF3D7_0832800 | rifin | 58269 | 0.069067168 |
| PF3D7_0500500 | rifin | 61281 | 0.07263734 |
| PF3D7_0901000 | rifin | 74182 | 0.087929099 |

**Extended Data Table 3:**
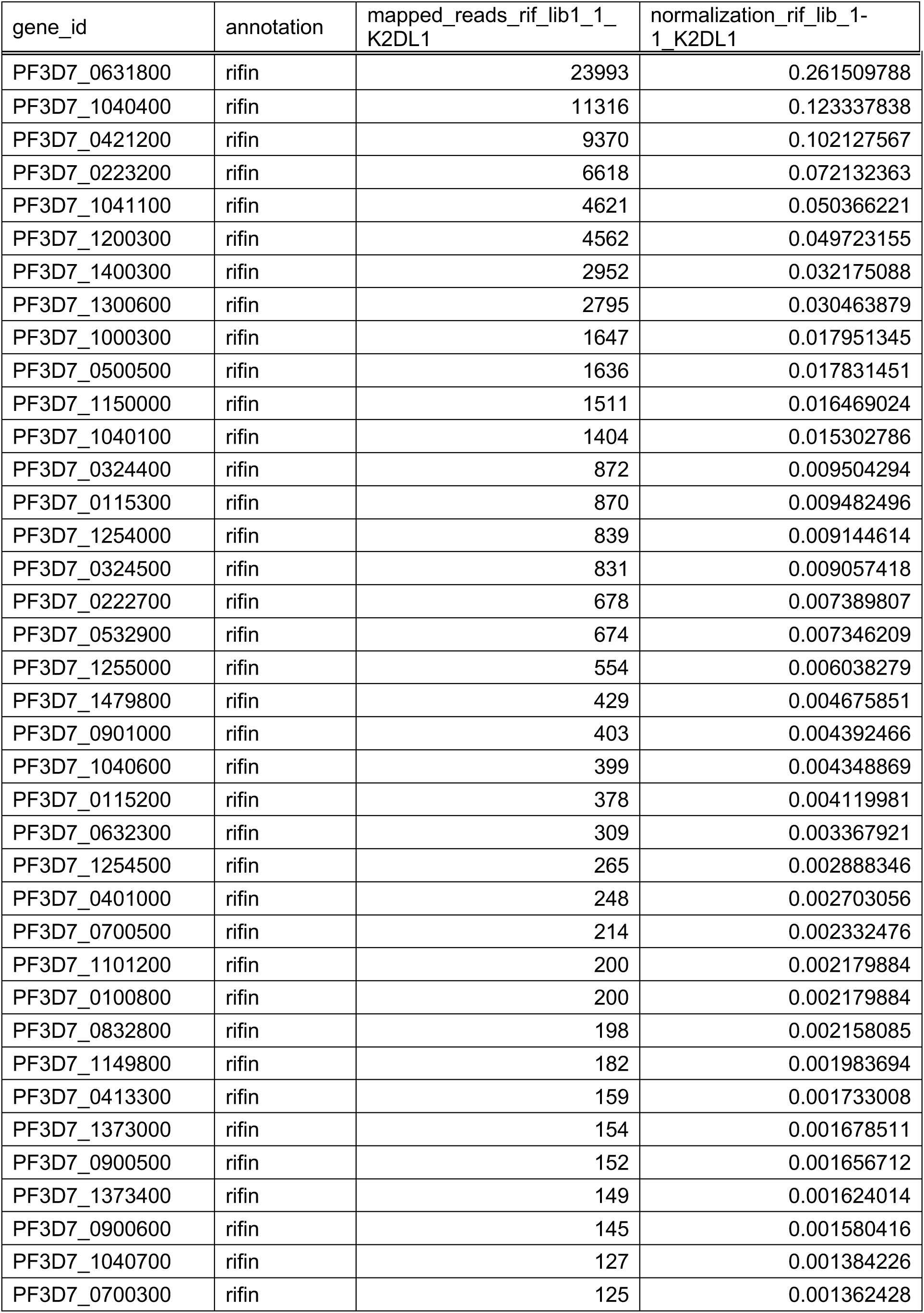
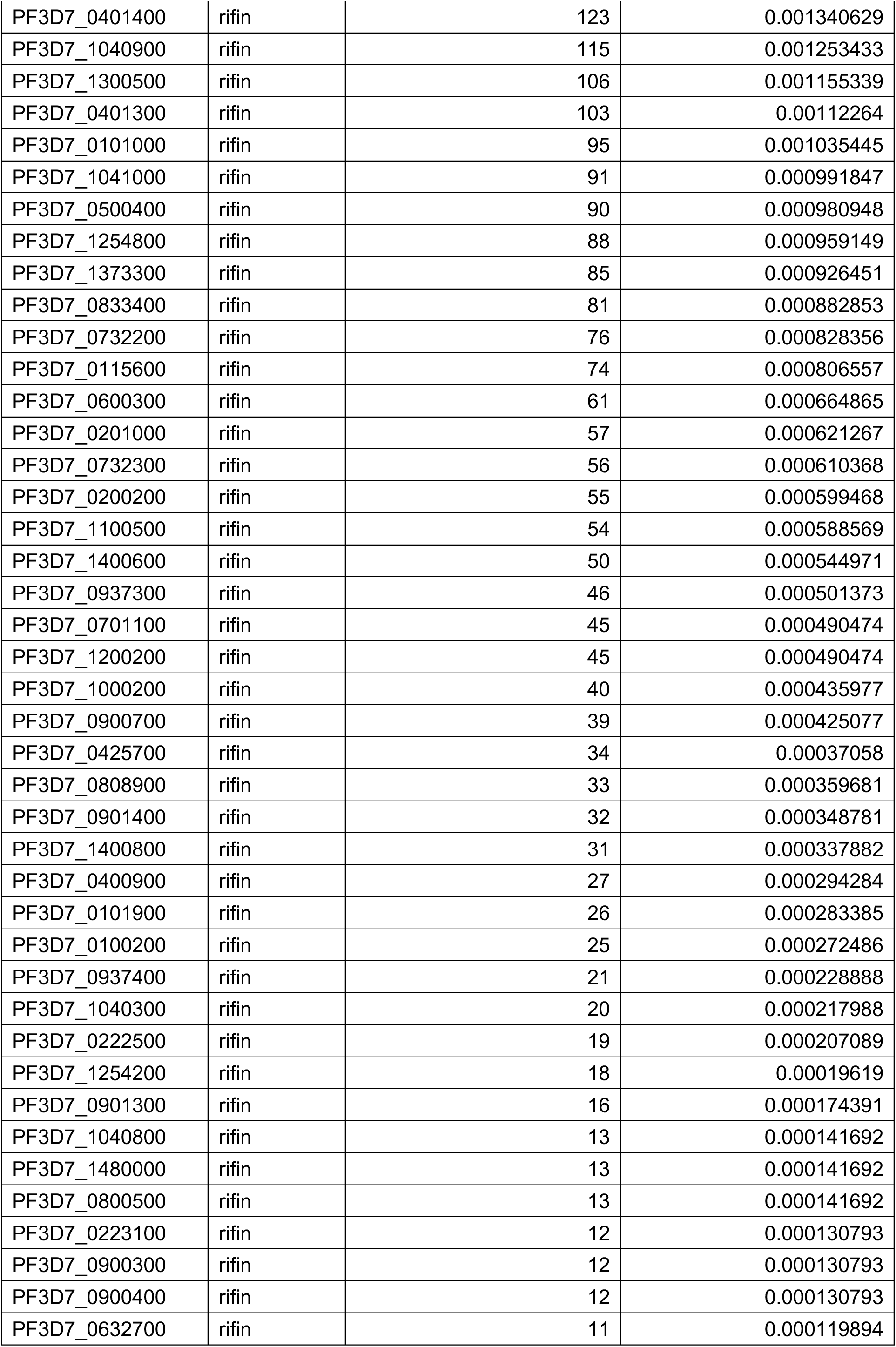
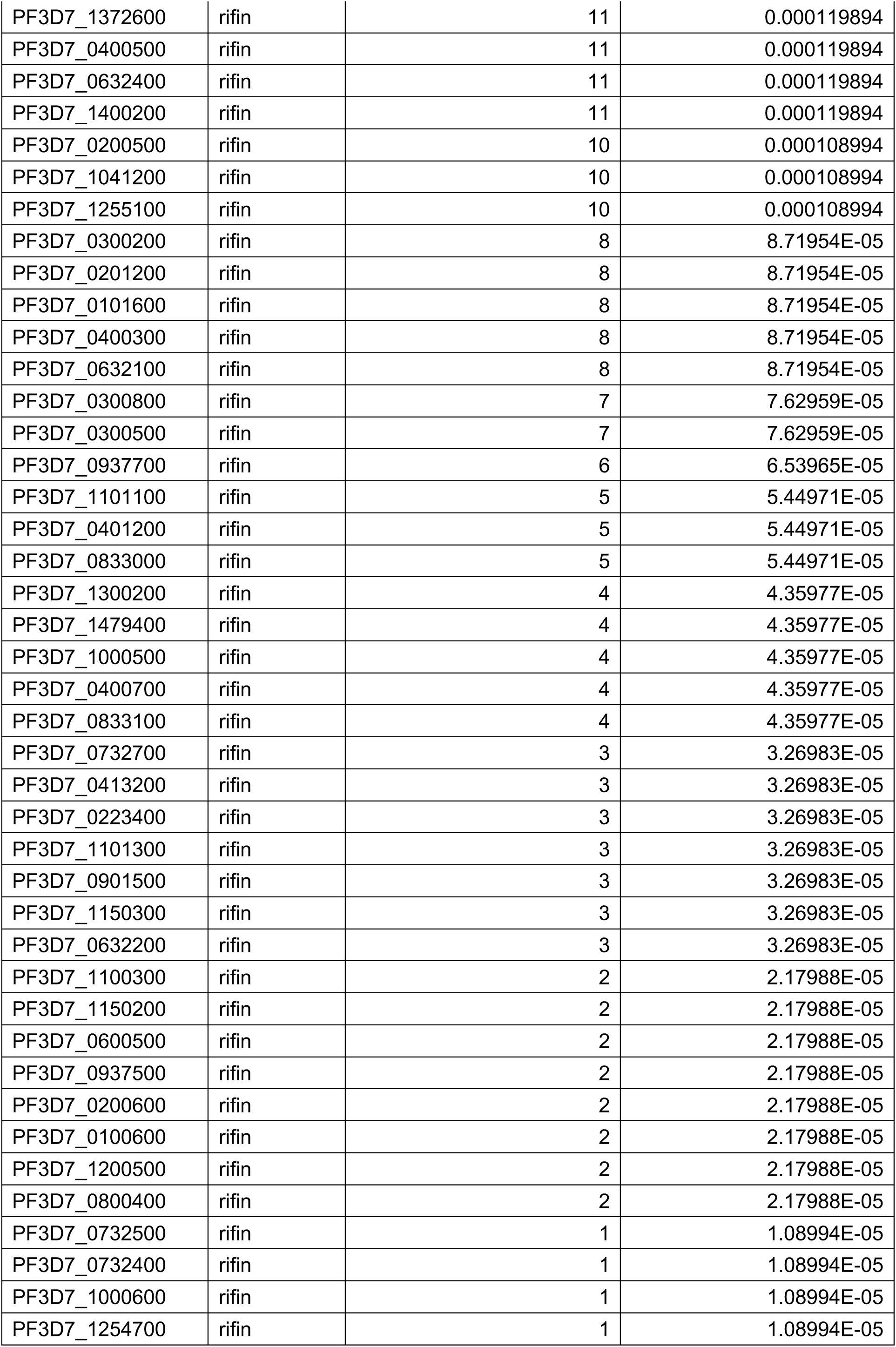
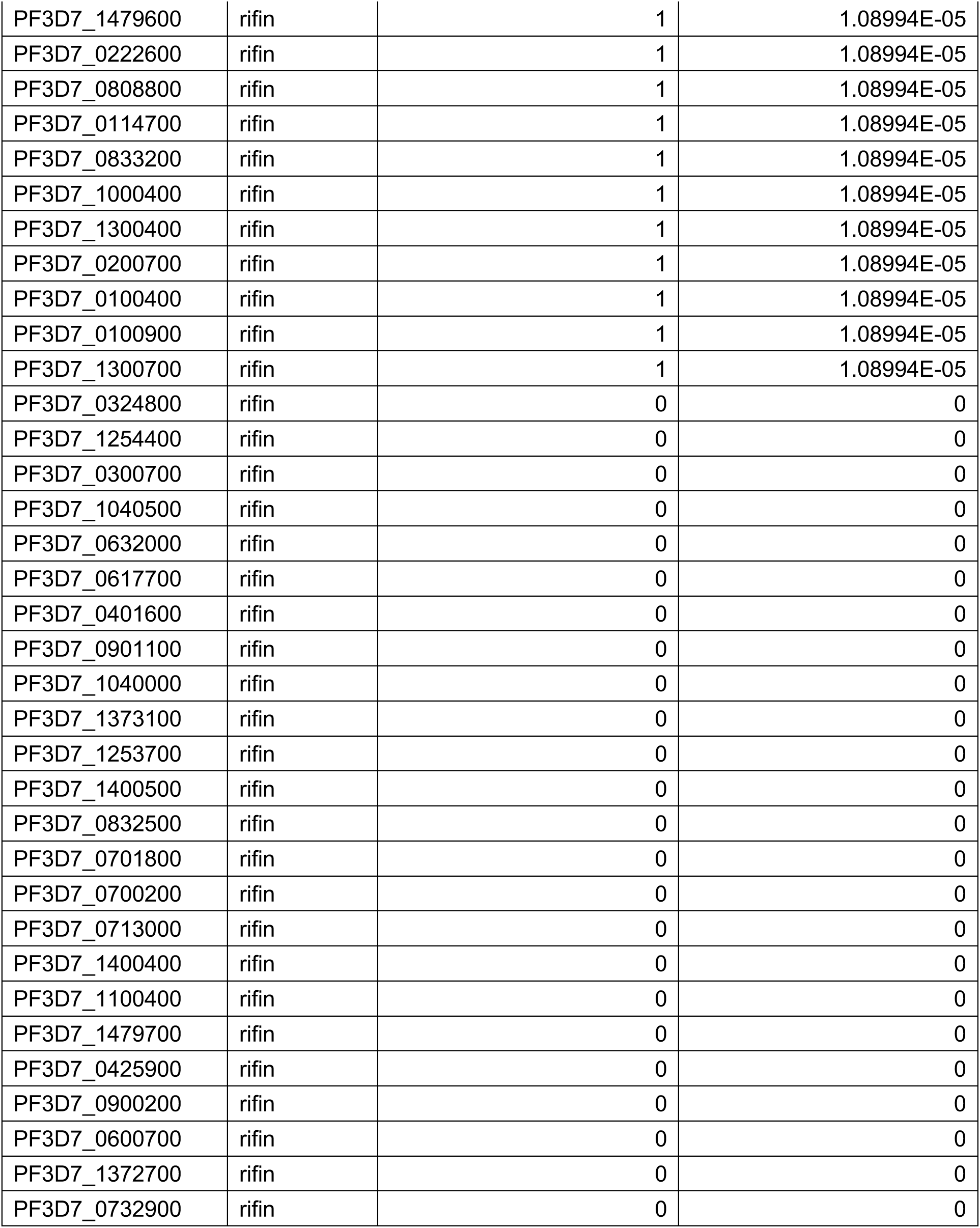
Summary of mapping results after screening of rif-lib1 with KIR2DL1.

**Extended Data Table 4:** Summary of mapping results after screening of rif-lib2 with KIR2DL1.

| gene_id | annotation | mapped_reads_rif_lib1_2_K2DL1 | normalization_rif_lib1_2_K2DL1 |
| --- | --- | --- | --- |
| PF3D7_1040400 | rifin | 22844 | 0.457639682 |
| PF3D7_0223200 | rifin | 2937 | 0.058837671 |
| PF3D7_1400300 | rifin | 2376 | 0.047599014 |
| PF3D7_0631800 | rifin | 1692 | 0.033896268 |
| PF3D7_1300600 | rifin | 1616 | 0.03237374 |
| PF3D7_1150000 | rifin | 1550 | 0.031051546 |
| PF3D7_1041100 | rifin | 1253 | 0.025101669 |
| PF3D7_1200300 | rifin | 1120 | 0.022437246 |
| PF3D7_0732300 | rifin | 1097 | 0.021976481 |
| PF3D7_0901000 | rifin | 952 | 0.019071659 |
| PF3D7_1000300 | rifin | 826 | 0.016547469 |
| PF3D7_0324500 | rifin | 822 | 0.016467336 |
| PF3D7_0500500 | rifin | 799 | 0.016006571 |
| PF3D7_0421200 | rifin | 756 | 0.015145141 |
| PF3D7_0324400 | rifin | 494 | 0.009896428 |
| PF3D7_0632300 | rifin | 436 | 0.008734499 |
| PF3D7_0832800 | rifin | 392 | 0.007853036 |
| PF3D7_1254500 | rifin | 392 | 0.007853036 |
| PF3D7_1254000 | rifin | 308 | 0.006170243 |
| PF3D7_1255000 | rifin | 308 | 0.006170243 |
| PF3D7_0900600 | rifin | 296 | 0.005929844 |
| PF3D7_1040100 | rifin | 259 | 0.005188613 |
| PF3D7_0532900 | rifin | 210 | 0.004206984 |
| PF3D7_0115300 | rifin | 202 | 0.004046718 |
| PF3D7_0700300 | rifin | 188 | 0.003766252 |
| PF3D7_0413300 | rifin | 155 | 0.003105155 |
| PF3D7_0401000 | rifin | 148 | 0.002964922 |
| PF3D7_1040700 | rifin | 125 | 0.002504157 |
| PF3D7_0101000 | rifin | 124 | 0.002484124 |
| PF3D7_0100800 | rifin | 122 | 0.002444057 |
| PF3D7_0401400 | rifin | 101 | 0.002023359 |
| PF3D7_1040600 | rifin | 93 | 0.001863093 |
| PF3D7_0500400 | rifin | 87 | 0.001742893 |
| PF3D7_1479800 | rifin | 86 | 0.00172286 |
| PF3D7_1373400 | rifin | 83 | 0.00166276 |
| PF3D7_1101200 | rifin | 76 | 0.001522527 |
| PF3D7_0900500 | rifin | 71 | 0.001422361 |
| PF3D7_1040900 | rifin | 69 | 0.001382295 |
| PF3D7_0201000 | rifin | 64 | 0.001282128 |
| PF3D7_0732200 | rifin | 61 | 0.001222029 |
| PF3D7_1041000 | rifin | 59 | 0.001181962 |
| PF3D7_1100500 | rifin | 58 | 0.001161929 |
| PF3D7_1254800 | rifin | 53 | 0.001061763 |
| PF3D7_1149800 | rifin | 53 | 0.001061763 |
| PF3D7_1400800 | rifin | 52 | 0.001041729 |
| PF3D7_0115600 | rifin | 51 | 0.001021696 |
| PF3D7_1300500 | rifin | 51 | 0.001021696 |
| PF3D7_0222700 | rifin | 51 | 0.001021696 |
| PF3D7_0600300 | rifin | 48 | 0.000961596 |
| PF3D7_0700500 | rifin | 48 | 0.000961596 |
| PF3D7_0100200 | rifin | 37 | 0.00074123 |
| PF3D7_0937300 | rifin | 37 | 0.00074123 |
| PF3D7_0808900 | rifin | 35 | 0.000701164 |
| PF3D7_1000200 | rifin | 30 | 0.000600998 |
| PF3D7_1254200 | rifin | 29 | 0.000580964 |
| PF3D7_0115200 | rifin | 29 | 0.000580964 |
| PF3D7_0701100 | rifin | 27 | 0.000540898 |
| PF3D7_0632700 | rifin | 24 | 0.000480798 |
| PF3D7_0200200 | rifin | 24 | 0.000480798 |
| PF3D7_0101900 | rifin | 20 | 0.000400665 |
| PF3D7_1200200 | rifin | 20 | 0.000400665 |
| PF3D7_0400900 | rifin | 19 | 0.000380632 |
| PF3D7_1040300 | rifin | 19 | 0.000380632 |
| PF3D7_0222500 | rifin | 18 | 0.000360599 |
| PF3D7_1480000 | rifin | 18 | 0.000360599 |
| PF3D7_0901300 | rifin | 16 | 0.000320532 |
| PF3D7_0937400 | rifin | 14 | 0.000280466 |
| PF3D7_1255100 | rifin | 12 | 0.000240399 |
| PF3D7_0833400 | rifin | 11 | 0.000220366 |
| PF3D7_0400300 | rifin | 11 | 0.000220366 |
| PF3D7_0937700 | rifin | 10 | 0.000200333 |
| PF3D7_1373300 | rifin | 10 | 0.000200333 |
| PF3D7_1400600 | rifin | 9 | 0.000180299 |
| PF3D7_0901400 | rifin | 8 | 0.000160266 |
| PF3D7_0300500 | rifin | 7 | 0.000140233 |
| PF3D7_0425700 | rifin | 6 | 0.0001202 |
| PF3D7_0900300 | rifin | 6 | 0.0001202 |
| PF3D7_0201200 | rifin | 6 | 0.0001202 |
| PF3D7_0900400 | rifin | 6 | 0.0001202 |
| PF3D7_1000500 | rifin | 6 | 0.0001202 |
| PF3D7_0833000 | rifin | 6 | 0.0001202 |
| PF3D7_0101600 | rifin | 6 | 0.0001202 |
| PF3D7_0900700 | rifin | 5 | 0.000100166 |
| PF3D7_0732400 | rifin | 5 | 0.000100166 |
| PF3D7_0632200 | rifin | 5 | 0.000100166 |
| PF3D7_0200500 | rifin | 4 | 8.0133E-05 |
| PF3D7_0401200 | rifin | 4 | 8.0133E-05 |
| PF3D7_1372600 | rifin | 4 | 8.0133E-05 |
| PF3D7_0100600 | rifin | 4 | 8.0133E-05 |
| PF3D7_0800400 | rifin | 4 | 8.0133E-05 |
| PF3D7_0632100 | rifin | 4 | 8.0133E-05 |
| PF3D7_1040800 | rifin | 3 | 6.00998E-05 |
| PF3D7_0732500 | rifin | 3 | 6.00998E-05 |
| PF3D7_0300800 | rifin | 3 | 6.00998E-05 |
| PF3D7_0600700 | rifin | 3 | 6.00998E-05 |
| PF3D7_0400700 | rifin | 3 | 6.00998E-05 |
| PF3D7_1254700 | rifin | 2 | 4.00665E-05 |
| PF3D7_1040000 | rifin | 2 | 4.00665E-05 |
| PF3D7_0901500 | rifin | 2 | 4.00665E-05 |
| PF3D7_0400500 | rifin | 2 | 4.00665E-05 |
| PF3D7_1373000 | rifin | 2 | 4.00665E-05 |
| PF3D7_0632400 | rifin | 2 | 4.00665E-05 |
| PF3D7_1000600 | rifin | 1 | 2.00333E-05 |
| PF3D7_0223100 | rifin | 1 | 2.00333E-05 |
| PF3D7_0632000 | rifin | 1 | 2.00333E-05 |
| PF3D7_0732700 | rifin | 1 | 2.00333E-05 |
| PF3D7_1101100 | rifin | 1 | 2.00333E-05 |
| PF3D7_0413200 | rifin | 1 | 2.00333E-05 |
| PF3D7_0223400 | rifin | 1 | 2.00333E-05 |
| PF3D7_0300200 | rifin | 1 | 2.00333E-05 |
| PF3D7_1101300 | rifin | 1 | 2.00333E-05 |
| PF3D7_1400500 | rifin | 1 | 2.00333E-05 |
| PF3D7_0401300 | rifin | 1 | 2.00333E-05 |
| PF3D7_0808800 | rifin | 1 | 2.00333E-05 |
| PF3D7_1479400 | rifin | 1 | 2.00333E-05 |
| PF3D7_1041200 | rifin | 1 | 2.00333E-05 |
| PF3D7_0600500 | rifin | 1 | 2.00333E-05 |
| PF3D7_0833200 | rifin | 1 | 2.00333E-05 |
| PF3D7_0937500 | rifin | 1 | 2.00333E-05 |
| PF3D7_0732900 | rifin | 1 | 2.00333E-05 |
| PF3D7_0800500 | rifin | 1 | 2.00333E-05 |
| PF3D7_1200500 | rifin | 1 | 2.00333E-05 |
| PF3D7_0100900 | rifin | 1 | 2.00333E-05 |
| PF3D7_1300700 | rifin | 1 | 2.00333E-05 |
| PF3D7_0324800 | rifin | 0 | 0 |
| PF3D7_1254400 | rifin | 0 | 0 |
| PF3D7_0300700 | rifin | 0 | 0 |
| PF3D7_1040500 | rifin | 0 | 0 |
| PF3D7_0617700 | rifin | 0 | 0 |
| PF3D7_1300200 | rifin | 0 | 0 |
| PF3D7_0401600 | rifin | 0 | 0 |
| PF3D7_0901100 | rifin | 0 | 0 |
| PF3D7_1479600 | rifin | 0 | 0 |
| PF3D7_1373100 | rifin | 0 | 0 |
| PF3D7_1253700 | rifin | 0 | 0 |
| PF3D7_0222600 | rifin | 0 | 0 |
| PF3D7_1100300 | rifin | 0 | 0 |
| PF3D7_1150300 | rifin | 0 | 0 |
| PF3D7_0832500 | rifin | 0 | 0 |
| PF3D7_0701800 | rifin | 0 | 0 |
| PF3D7_0700200 | rifin | 0 | 0 |
| PF3D7_1150200 | rifin | 0 | 0 |
| PF3D7_0713000 | rifin | 0 | 0 |
| PF3D7_1400400 | rifin | 0 | 0 |
| PF3D7_1100400 | rifin | 0 | 0 |
| PF3D7_1479700 | rifin | 0 | 0 |
| PF3D7_0114700 | rifin | 0 | 0 |
| PF3D7_1000400 | rifin | 0 | 0 |
| PF3D7_1300400 | rifin | 0 | 0 |
| PF3D7_0200700 | rifin | 0 | 0 |
| PF3D7_0425900 | rifin | 0 | 0 |
| PF3D7_0900200 | rifin | 0 | 0 |
| PF3D7_1400200 | rifin | 0 | 0 |
| PF3D7_1372700 | rifin | 0 | 0 |
| PF3D7_0200600 | rifin | 0 | 0 |
| PF3D7_0100400 | rifin | 0 | 0 |
| PF3D7_0833100 | rifin | 0 | 0 |

**Extended Data Table 5:** The list of KIR2DL1-binding RIFINs identified from rif-lib1.

| gene_id | annotation | normalization_rif<br>lib_1-1_before | normalization_rif<br>lib_1-1_K2DL1 | fold_<br>enrichment |
| --- | --- | --- | --- | --- |
| PF3D7_1000600 | rifin | 0 | 1.08994E-05 | #DIV/0! |
| PF3D7_0401600 | rifin | 0 | 0 | #DIV/0! |
| PF3D7_0901100 | rifin | 0 | 0 | #DIV/0! |
| PF3D7_1253700 | rifin | 0 | 0 | #DIV/0! |
| PF3D7_0832500 | rifin | 0 | 0 | #DIV/0! |
| PF3D7_0701800 | rifin | 0 | 0 | #DIV/0! |
| PF3D7_0700200 | rifin | 0 | 0 | #DIV/0! |
| PF3D7_1040400 | rifin | 0.002494771 | 0.123337838 | 49.43853236 |
| PF3D7_0631800 | rifin | 0.008400558 | 0.261509788 | 31.13005072 |
| PF3D7_0421200 | rifin | 0.004949706 | 0.102127567 | 20.63305631 |
| PF3D7_0632300 | rifin | 0.000224081 | 0.003367921 | 15.02990801 |
| PF3D7_0222700 | rifin | 0.000587591 | 0.007389807 | 12.57644927 |
| PF3D7_1300600 | rifin | 0.002708894 | 0.030463879 | 11.24587545 |
| PF3D7_0900500 | rifin | 0.000154367 | 0.001656712 | 10.73228633 |
| PF3D7_1041100 | rifin | 0.004974604 | 0.050366221 | 10.12466907 |
| PF3D7_0115300 | rifin | 0.001404243 | 0.009482496 | 6.75274735 |
| PF3D7_0223200 | rifin | 0.013673937 | 0.072132363 | 5.27517154 |
| PF3D7_1101200 | rifin | 0.000418285 | 0.002179884 | 5.211479892 |
| PF3D7_0223100 | rifin | 3.48571E-05 | 0.000130793 | 3.752265522 |
| PF3D7_0400900 | rifin | 8.46529E-05 | 0.000294284 | 3.476363646 |
| PF3D7_1041000 | rifin | 0.000298775 | 0.000991847 | 3.319712691 |
| PF3D7_1200300 | rifin | 0.01579026 | 0.049723155 | 3.148976327 |
| PF3D7_0413300 | rifin | 0.00059755 | 0.001733008 | 2.90018856 |
| PF3D7_1400300 | rifin | 0.011094512 | 0.032175088 | 2.900090318 |
| PF3D7_1255000 | rifin | 0.002923016 | 0.006038279 | 2.065770258 |
| PF3D7_0937400 | rifin | 0.00011453 | 0.000228888 | 1.998489246 |
| PF3D7_1150300 | rifin | 1.99183E-05 | 3.26983E-05 | 1.641616166 |
| PF3D7_0700500 | rifin | 0.001454038 | 0.002332476 | 1.604136345 |
| PF3D7_0833000 | rifin | 3.48571E-05 | 5.44971E-05 | 1.563443968 |
| PF3D7_0324500 | rifin | 0.006075092 | 0.009057418 | 1.49091042 |
| PF3D7_0600500 | rifin | 1.49388E-05 | 2.17988E-05 | 1.45921437 |
| PF3D7_0100800 | rifin | 0.001583508 | 0.002179884 | 1.37661733 |
| PF3D7_0900700 | rifin | 0.000318693 | 0.000425077 | 1.333813135 |
| PF3D7_0413200 | rifin | 2.48979E-05 | 3.26983E-05 | 1.313292933 |
| PF3D7_1400800 | rifin | 0.000273877 | 0.000337882 | 1.233699422 |
| PF3D7_1000200 | rifin | 0.000408326 | 0.000435977 | 1.067717832 |
| PF3D7_0115200 | rifin | 0.003899014 | 0.004119981 | 1.056672475 |
| PF3D7_1040800 | rifin | 0.000134449 | 0.000141692 | 1.053877045 |
| PF3D7_1254800 | rifin | 0.000911264 | 0.000959149 | 1.05254807 |
| PF3D7_0300200 | rifin | 8.96325E-05 | 8.71954E-05 | 0.97280958 |
| PF3D7_0425700 | rifin | 0.000388408 | 0.00037058 | 0.954101703 |
| PF3D7_1150200 | rifin | 2.48979E-05 | 2.17988E-05 | 0.875528622 |
| PF3D7_0900300 | rifin | 0.000154367 | 0.000130793 | 0.847285763 |
| PF3D7_0115600 | rifin | 0.000985958 | 0.000806557 | 0.818044419 |
| PF3D7_0701100 | rifin | 0.000632407 | 0.000490474 | 0.775566693 |
| PF3D7_0732200 | rifin | 0.001085549 | 0.000828356 | 0.763075404 |
| PF3D7_1150000 | rifin | 0.022428045 | 0.016469024 | 0.734304922 |
| PF3D7_0100400 | rifin | 1.49388E-05 | 1.08994E-05 | 0.729607185 |
| PF3D7_1300700 | rifin | 1.49388E-05 | 1.08994E-05 | 0.729607185 |
| PF3D7_1400200 | rifin | 0.000179265 | 0.000119894 | 0.668806586 |
| PF3D7_0937700 | rifin | 9.95917E-05 | 6.53965E-05 | 0.656646466 |
| PF3D7_0632400 | rifin | 0.000189224 | 0.000119894 | 0.63360624 |
| PF3D7_1041200 | rifin | 0.000184245 | 0.000108994 | 0.591573393 |
| PF3D7_1040700 | rifin | 0.002474853 | 0.001384226 | 0.559316574 |
| PF3D7_0901300 | rifin | 0.00035355 | 0.000174391 | 0.493255562 |
| PF3D7_0401300 | rifin | 0.002290609 | 0.00112264 | 0.490105696 |
| PF3D7_1101100 | rifin | 0.00011453 | 5.44971E-05 | 0.475830773 |
| PF3D7_0324400 | rifin | 0.01997809 | 0.009504294 | 0.475735891 |
| PF3D7_0101900 | rifin | 0.000642366 | 0.000283385 | 0.441157833 |
| PF3D7_1000400 | rifin | 2.48979E-05 | 1.08994E-05 | 0.437764311 |
| PF3D7_0100600 | rifin | 4.97958E-05 | 2.17988E-05 | 0.437764311 |
| PF3D7_1000300 | rifin | 0.042749726 | 0.017951345 | 0.419917193 |
| PF3D7_0300800 | rifin | 0.000184245 | 7.62959E-05 | 0.414101375 |
| PF3D7_1480000 | rifin | 0.000348571 | 0.000141692 | 0.406495432 |
| PF3D7_0732700 | rifin | 8.46529E-05 | 3.26983E-05 | 0.386262627 |
| PF3D7_0937300 | rifin | 0.001299671 | 0.000501373 | 0.385769316 |
| PF3D7_0808900 | rifin | 0.000936162 | 0.000359681 | 0.384208039 |
| PF3D7_0400700 | rifin | 0.00011453 | 4.35977E-05 | 0.380664618 |
| PF3D7_1255100 | rifin | 0.000293795 | 0.000108994 | 0.370986704 |
| PF3D7_0500400 | rifin | 0.002644159 | 0.000980948 | 0.370986704 |
| PF3D7_1254200 | rifin | 0.000552734 | 0.00019619 | 0.354944036 |
| PF3D7_0901500 | rifin | 9.95917E-05 | 3.26983E-05 | 0.328323233 |
| PF3D7_0900400 | rifin | 0.000403346 | 0.000130793 | 0.32426986 |
| PF3D7_0400500 | rifin | 0.000373469 | 0.000119894 | 0.321027161 |
| PF3D7_0200200 | rifin | 0.001872323 | 0.000599468 | 0.320173366 |
| PF3D7_1479600 | rifin | 3.48571E-05 | 1.08994E-05 | 0.312688794 |
| PF3D7_0800400 | rifin | 6.97142E-05 | 2.17988E-05 | 0.312688794 |
| PF3D7_1149800 | rifin | 0.00645354 | 0.001983694 | 0.307380805 |
| PF3D7_0400300 | rifin | 0.000288816 | 8.71954E-05 | 0.301906421 |
| PF3D7_1479800 | rifin | 0.015690668 | 0.004675851 | 0.298002046 |
| PF3D7_0800500 | rifin | 0.00048302 | 0.000141692 | 0.293347219 |
| PF3D7_1479400 | rifin | 0.000149388 | 4.35977E-05 | 0.291842874 |
| PF3D7_0632100 | rifin | 0.000308734 | 8.71954E-05 | 0.282428588 |
| PF3D7_0401400 | rifin | 0.0047804 | 0.001340629 | 0.280442762 |
| PF3D7_0808800 | rifin | 3.98367E-05 | 1.08994E-05 | 0.273602694 |
| PF3D7_0100900 | rifin | 3.98367E-05 | 1.08994E-05 | 0.273602694 |
| PF3D7_0223400 | rifin | 0.00011951 | 3.26983E-05 | 0.273602694 |
| PF3D7_1200200 | rifin | 0.002046609 | 0.000490474 | 0.239651995 |
| PF3D7_0401200 | rifin | 0.000229061 | 5.44971E-05 | 0.237915386 |
| PF3D7_1254000 | rifin | 0.03983169 | 0.009144614 | 0.229581358 |
| PF3D7_0600300 | rifin | 0.002967832 | 0.000664865 | 0.224023683 |
| PF3D7_1373000 | rifin | 0.007917538 | 0.001678511 | 0.211999069 |
| PF3D7_0300500 | rifin | 0.000378448 | 7.62959E-05 | 0.201601985 |
| PF3D7_0901400 | rifin | 0.001737875 | 0.000348781 | 0.20069424 |
| PF3D7_0732500 | rifin | 5.47754E-05 | 1.08994E-05 | 0.198983778 |
| PF3D7_0833200 | rifin | 5.47754E-05 | 1.08994E-05 | 0.198983778 |
| PF3D7_0100200 | rifin | 0.001389304 | 0.000272486 | 0.196130964 |
| PF3D7_1300200 | rifin | 0.000224081 | 4.35977E-05 | 0.194561916 |
| PF3D7_0833100 | rifin | 0.000224081 | 4.35977E-05 | 0.194561916 |
| PF3D7_0632700 | rifin | 0.000632407 | 0.000119894 | 0.189582969 |
| PF3D7_0200700 | rifin | 5.9755E-05 | 1.08994E-05 | 0.182401796 |
| PF3D7_1101300 | rifin | 0.000184245 | 3.26983E-05 | 0.177472018 |
| PF3D7_0201000 | rifin | 0.003545464 | 0.000621267 | 0.175228692 |
| PF3D7_0200500 | rifin | 0.000642366 | 0.000108994 | 0.16967609 |
| PF3D7_1400600 | rifin | 0.003336321 | 0.000544971 | 0.163344892 |
| PF3D7_1300500 | rifin | 0.007115825 | 0.001155339 | 0.162361851 |
| PF3D7_1040100 | rifin | 0.100492979 | 0.015302786 | 0.152277165 |
| PF3D7_0200600 | rifin | 0.000144408 | 2.17988E-05 | 0.150953211 |
| PF3D7_1200500 | rifin | 0.000144408 | 2.17988E-05 | 0.150953211 |
| PF3D7_1373300 | rifin | 0.006169704 | 0.000926451 | 0.150161285 |
| PF3D7_1372600 | rifin | 0.000811672 | 0.000119894 | 0.147711884 |
| PF3D7_0732400 | rifin | 7.46938E-05 | 1.08994E-05 | 0.145921437 |
| PF3D7_0500500 | rifin | 0.122701922 | 0.017831451 | 0.145323325 |
| PF3D7_0901000 | rifin | 0.030250971 | 0.004392466 | 0.145200837 |
| PF3D7_1040600 | rifin | 0.030539787 | 0.004348869 | 0.142400098 |
| PF3D7_0732300 | rifin | 0.00430236 | 0.000610368 | 0.141868064 |
| PF3D7_0900600 | rifin | 0.011467981 | 0.001580416 | 0.13781117 |
| PF3D7_1000500 | rifin | 0.000338612 | 4.35977E-05 | 0.128754209 |
| PF3D7_0101000 | rifin | 0.008226272 | 0.001035445 | 0.125870489 |
| PF3D7_0401000 | rifin | 0.021556618 | 0.002703056 | 0.125393335 |
| PF3D7_0632200 | rifin | 0.000298775 | 3.26983E-05 | 0.109441078 |
| PF3D7_0101600 | rifin | 0.000801713 | 8.71954E-05 | 0.108761319 |
| PF3D7_0222500 | rifin | 0.002086446 | 0.000207089 | 0.099254438 |
| PF3D7_0532900 | rifin | 0.074853102 | 0.007346209 | 0.09814168 |
| PF3D7_1100500 | rifin | 0.00645354 | 0.000588569 | 0.091200898 |
| PF3D7_1040900 | rifin | 0.013783488 | 0.001253433 | 0.090937312 |
| PF3D7_1100300 | rifin | 0.000263918 | 2.17988E-05 | 0.08259704 |
| PF3D7_1040300 | rifin | 0.002664077 | 0.000217988 | 0.081825105 |
| PF3D7_0201200 | rifin | 0.001100488 | 8.71954E-05 | 0.079233359 |
| PF3D7_0937500 | rifin | 0.000278857 | 2.17988E-05 | 0.078172198 |
| PF3D7_1254700 | rifin | 0.000154367 | 1.08994E-05 | 0.070607147 |
| PF3D7_0700300 | rifin | 0.020560701 | 0.001362428 | 0.06626367 |
| PF3D7_0833400 | rifin | 0.013355244 | 0.000882853 | 0.066105349 |
| PF3D7_1373400 | rifin | 0.027845832 | 0.001624014 | 0.058321604 |
| PF3D7_1254500 | rifin | 0.054073299 | 0.002888346 | 0.053415389 |
| PF3D7_0114700 | rifin | 0.000214122 | 1.08994E-05 | 0.050902827 |
| PF3D7_1300400 | rifin | 0.000268898 | 1.08994E-05 | 0.040533732 |
| PF3D7_0832800 | rifin | 0.08073897 | 0.002158085 | 0.026729164 |
| PF3D7_0222600 | rifin | 0.000413305 | 1.08994E-05 | 0.026371344 |
| PF3D7_0632000 | rifin | 4.97958E-06 | 0 | 0 |
| PF3D7_1100400 | rifin | 4.97958E-06 | 0 | 0 |
| PF3D7_0732900 | rifin | 4.97958E-06 | 0 | 0 |
| PF3D7_0617700 | rifin | 9.95917E-06 | 0 | 0 |
| PF3D7_1400400 | rifin | 1.49388E-05 | 0 | 0 |
| PF3D7_0600700 | rifin | 1.49388E-05 | 0 | 0 |
| PF3D7_0425900 | rifin | 1.99183E-05 | 0 | 0 |
| PF3D7_1254400 | rifin | 2.98775E-05 | 0 | 0 |
| PF3D7_0300700 | rifin | 2.98775E-05 | 0 | 0 |
| PF3D7_0713000 | rifin | 4.48163E-05 | 0 | 0 |
| PF3D7_1479700 | rifin | 4.48163E-05 | 0 | 0 |
| PF3D7_1040500 | rifin | 5.9755E-05 | 0 | 0 |
| PF3D7_0324800 | rifin | 6.97142E-05 | 0 | 0 |
| PF3D7_0900200 | rifin | 7.46938E-05 | 0 | 0 |
| PF3D7_1040000 | rifin | 0.000149388 | 0 | 0 |
| PF3D7_1400500 | rifin | 0.000378448 | 0 | 0 |
| PF3D7_1373100 | rifin | 0.000428244 | 0 | 0 |
| PF3D7_1372700 | rifin | 0.000806693 | 0 | 0 |

**Extended Data Table 6:** The list of KIR2DL1-binding RIFINs identified from rif-lib2.

| gene_id | annotation | normalization_rif_lib<br>1-2_before | normalization_rif_lib<br>1_2_K2DL1 | fold_enrichment |
| --- | --- | --- | --- | --- |
| PF3D7_0401600 | rifin | 0 | 0 | #DIV/0! |
| PF3D7_1253700 | rifin | 0 | 0 | #DIV/0! |
| PF3D7_0832500 | rifin | 0 | 0 | #DIV/0! |
| PF3D7_0700200 | rifin | 0 | 0 | #DIV/0! |
| PF3D7_0632300 | rifin | 0.000267881 | 0.008734499 | 32.60584712 |
| PF3D7_0631800 | rifin | 0.001308589 | 0.033896268 | 25.90291993 |
| PF3D7_0421200 | rifin | 0.000613994 | 0.015145141 | 24.66661036 |
| PF3D7_1040400 | rifin | 0.018966239 | 0.457639682 | 24.12917449 |
| PF3D7_1041100 | rifin | 0.002332702 | 0.025101669 | 10.76077163 |
| PF3D7_1300600 | rifin | 0.00345164 | 0.03237374 | 9.379235135 |
| PF3D7_1041000 | rifin | 0.000126829 | 0.001181962 | 9.319351057 |
| PF3D7_0400900 | rifin | 5.21539E-05 | 0.000380632 | 7.298243721 |
| PF3D7_0413300 | rifin | 0.000574878 | 0.003105155 | 5.40141315 |
| PF3D7_1400800 | rifin | 0.000214542 | 0.001041729 | 4.855592217 |
| PF3D7_0900500 | rifin | 0.000302256 | 0.001422361 | 4.705823196 |
| PF3D7_0223200 | rifin | 0.013709363 | 0.058837671 | 4.29178736 |
| PF3D7_1000600 | rifin | 4.74126E-06 | 2.00333E-05 | 4.225298996 |
| PF3D7_0732300 | rifin | 0.005331551 | 0.021976481 | 4.121967985 |
| PF3D7_0937700 | rifin | 5.21539E-05 | 0.000200333 | 3.841180906 |
| PF3D7_0222700 | rifin | 0.000295144 | 0.001021696 | 3.461690744 |
| PF3D7_0324500 | rifin | 0.004830162 | 0.016467336 | 3.409271926 |
| PF3D7_0632000 | rifin | 5.92658E-06 | 2.00333E-05 | 3.380239197 |
| PF3D7_0732900 | rifin | 5.92658E-06 | 2.00333E-05 | 3.380239197 |
| PF3D7_0115300 | rifin | 0.001299106 | 0.004046718 | 3.11500145 |
| PF3D7_1400300 | rifin | 0.017406363 | 0.047599014 | 2.73457553 |
| PF3D7_1254200 | rifin | 0.00022521 | 0.000580964 | 2.579656229 |
| PF3D7_1101200 | rifin | 0.000627032 | 0.001522527 | 2.42814914 |
| PF3D7_0632700 | rifin | 0.00022521 | 0.000480798 | 2.134887914 |
| PF3D7_1480000 | rifin | 0.000186095 | 0.000360599 | 1.937716737 |
| PF3D7_1255000 | rifin | 0.003312958 | 0.006170243 | 1.862457375 |
| PF3D7_0808900 | rifin | 0.000382857 | 0.000701164 | 1.831398946 |
| PF3D7_0937400 | rifin | 0.000160018 | 0.000280466 | 1.752716621 |
| PF3D7_0800400 | rifin | 4.74126E-05 | 8.0133E-05 | 1.690119599 |
| PF3D7_1040700 | rifin | 0.001527872 | 0.002504157 | 1.638983319 |
| PF3D7_0833000 | rifin | 7.34896E-05 | 0.0001202 | 1.635599611 |
| PF3D7_0100800 | rifin | 0.001521946 | 0.002444057 | 1.605876877 |
| PF3D7_1150000 | rifin | 0.019390582 | 0.031051546 | 1.601372564 |
| PF3D7_1255100 | rifin | 0.00015172 | 0.000240399 | 1.584487124 |
| PF3D7_1254800 | rifin | 0.000676815 | 0.001061763 | 1.5687625 |
| PF3D7_0115600 | rifin | 0.000689854 | 0.001021696 | 1.481032638 |
| PF3D7_0808800 | rifin | 1.54091E-05 | 2.00333E-05 | 1.300091999 |
| PF3D7_0101900 | rifin | 0.000346112 | 0.000400665 | 1.157616163 |
| PF3D7_0900700 | rifin | 9.0084E-05 | 0.000100166 | 1.111920789 |
| PF3D7_0400300 | rifin | 0.000200318 | 0.000220366 | 1.100077845 |
| PF3D7_1000200 | rifin | 0.000551172 | 0.000600998 | 1.090399741 |
| PF3D7_0732200 | rifin | 0.001149756 | 0.001222029 | 1.062858717 |
| PF3D7_0901500 | rifin | 4.03007E-05 | 4.00665E-05 | 0.994187999 |
| PF3D7_1300700 | rifin | 2.01504E-05 | 2.00333E-05 | 0.994187999 |
| PF3D7_0425700 | rifin | 0.000123273 | 0.0001202 | 0.975068999 |
| PF3D7_0100600 | rifin | 8.41574E-05 | 8.0133E-05 | 0.952180056 |
| PF3D7_0401400 | rifin | 0.002167943 | 0.002023359 | 0.933308253 |
| PF3D7_1000500 | rifin | 0.000142238 | 0.0001202 | 0.845059799 |
| PF3D7_0413200 | rifin | 2.48916E-05 | 2.00333E-05 | 0.804818856 |
| PF3D7_0700500 | rifin | 0.001231543 | 0.000961596 | 0.780805974 |
| PF3D7_0201000 | rifin | 0.001678407 | 0.001282128 | 0.763895864 |
| PF3D7_0937300 | rifin | 0.001040707 | 0.00074123 | 0.712237188 |
| PF3D7_0400700 | rifin | 9.12693E-05 | 6.00998E-05 | 0.658488155 |
| PF3D7_1040800 | rifin | 9.364E-05 | 6.00998E-05 | 0.641817569 |
| PF3D7_0732400 | rifin | 0.000161203 | 0.000100166 | 0.621367499 |
| PF3D7_0701100 | rifin | 0.000879504 | 0.000540898 | 0.615003088 |
| PF3D7_0300200 | rifin | 3.55595E-05 | 2.00333E-05 | 0.5633732 |
| PF3D7_0223100 | rifin | 3.67448E-05 | 2.00333E-05 | 0.54519987 |
| PF3D7_1200200 | rifin | 0.000752676 | 0.000400665 | 0.532321133 |
| PF3D7_0324400 | rifin | 0.018967424 | 0.009896428 | 0.521759206 |
| PF3D7_1300500 | rifin | 0.002023334 | 0.001021696 | 0.504956646 |
| PF3D7_0600300 | rifin | 0.001941547 | 0.000961596 | 0.495273142 |
| PF3D7_0732500 | rifin | 0.000124458 | 6.00998E-05 | 0.482891314 |
| PF3D7_1200300 | rifin | 0.048078781 | 0.022437246 | 0.46667668 |
| PF3D7_0900300 | rifin | 0.000258399 | 0.0001202 | 0.465170532 |
| PF3D7_0200200 | rifin | 0.001064414 | 0.000480798 | 0.451702343 |
| PF3D7_0600500 | rifin | 4.5042E-05 | 2.00333E-05 | 0.444768315 |
| PF3D7_0101000 | rifin | 0.005952656 | 0.002484124 | 0.417313481 |
| PF3D7_0101600 | rifin | 0.000289217 | 0.0001202 | 0.41560318 |
| PF3D7_0100200 | rifin | 0.001852649 | 0.00074123 | 0.400092291 |
| PF3D7_1000300 | rifin | 0.042532688 | 0.016547469 | 0.389052974 |
| PF3D7_0100900 | rifin | 5.21539E-05 | 2.00333E-05 | 0.384118091 |
| PF3D7_0200500 | rifin | 0.000213357 | 8.0133E-05 | 0.375582133 |
| PF3D7_0901300 | rifin | 0.00091862 | 0.000320532 | 0.348927917 |
| PF3D7_0300800 | rifin | 0.000175427 | 6.00998E-05 | 0.342591811 |
| PF3D7_0500400 | rifin | 0.005133603 | 0.001742893 | 0.339506823 |
| PF3D7_1040000 | rifin | 0.000123273 | 4.00665E-05 | 0.325023 |
| PF3D7_0900400 | rifin | 0.000398266 | 0.0001202 | 0.301807071 |
| PF3D7_1254000 | rifin | 0.020676649 | 0.006170243 | 0.29841598 |
| PF3D7_0632100 | rifin | 0.000271437 | 8.0133E-05 | 0.295217397 |
| PF3D7_0532900 | rifin | 0.015628389 | 0.004206984 | 0.269188559 |
| PF3D7_1372600 | rifin | 0.000303441 | 8.0133E-05 | 0.264081187 |
| PF3D7_0700300 | rifin | 0.014959871 | 0.003766252 | 0.25175698 |
| PF3D7_1479800 | rifin | 0.007382147 | 0.00172286 | 0.233381961 |
| PF3D7_0500500 | rifin | 0.07263734 | 0.016006571 | 0.220362846 |
| PF3D7_0901000 | rifin | 0.087929099 | 0.019071659 | 0.21689815 |
| PF3D7_0833200 | rifin | 9.24546E-05 | 2.00333E-05 | 0.216682 |
| PF3D7_1040900 | rifin | 0.006404261 | 0.001382295 | 0.215839815 |
| PF3D7_1149800 | rifin | 0.005036407 | 0.001061763 | 0.21081746 |
| PF3D7_0223400 | rifin | 9.60106E-05 | 2.00333E-05 | 0.208656741 |
| PF3D7_1100500 | rifin | 0.005971621 | 0.001161929 | 0.194575103 |
| PF3D7_0632400 | rifin | 0.00020743 | 4.00665E-05 | 0.193156526 |
| PF3D7_1254700 | rifin | 0.000208616 | 4.00665E-05 | 0.192059045 |
| PF3D7_0401200 | rifin | 0.000431455 | 8.0133E-05 | 0.185727428 |
| PF3D7_0401000 | rifin | 0.016205638 | 0.002964922 | 0.182956188 |
| PF3D7_0632200 | rifin | 0.00056184 | 0.000100166 | 0.178282658 |
| PF3D7_0201200 | rifin | 0.00075149 | 0.0001202 | 0.159948227 |
| PF3D7_1040300 | rifin | 0.002589915 | 0.000380632 | 0.146966922 |
| PF3D7_0600700 | rifin | 0.000413675 | 6.00998E-05 | 0.145282487 |
| PF3D7_0800500 | rifin | 0.000138682 | 2.00333E-05 | 0.144454667 |
| PF3D7_0300500 | rifin | 0.00105019 | 0.000140233 | 0.133530894 |
| PF3D7_1254500 | rifin | 0.059131851 | 0.007853036 | 0.132805517 |
| PF3D7_1200500 | rifin | 0.000163574 | 2.00333E-05 | 0.122472435 |
| PF3D7_0832800 | rifin | 0.069067168 | 0.007853036 | 0.113701433 |
| PF3D7_0222500 | rifin | 0.003220503 | 0.000360599 | 0.111969646 |
| PF3D7_1101300 | rifin | 0.000188465 | 2.00333E-05 | 0.10629683 |
| PF3D7_0401300 | rifin | 0.000188465 | 2.00333E-05 | 0.10629683 |
| PF3D7_0900600 | rifin | 0.060359838 | 0.005929844 | 0.098241541 |
| PF3D7_1101100 | rifin | 0.000210986 | 2.00333E-05 | 0.094950539 |
| PF3D7_1040100 | rifin | 0.054671507 | 0.005188613 | 0.09490525 |
| PF3D7_0901400 | rifin | 0.001760194 | 0.000160266 | 0.091050214 |
| PF3D7_1373400 | rifin | 0.021930714 | 0.00166276 | 0.075818791 |
| PF3D7_0400500 | rifin | 0.000552357 | 4.00665E-05 | 0.072537322 |
| PF3D7_0732700 | rifin | 0.000304626 | 2.00333E-05 | 0.065763409 |
| PF3D7_1040600 | rifin | 0.033878697 | 0.001863093 | 0.054993046 |
| PF3D7_1400600 | rifin | 0.003551206 | 0.000180299 | 0.050771283 |
| PF3D7_1041200 | rifin | 0.000440937 | 2.00333E-05 | 0.045433323 |
| PF3D7_1400500 | rifin | 0.000459903 | 2.00333E-05 | 0.043559783 |
| PF3D7_0115200 | rifin | 0.013640615 | 0.000580964 | 0.042590779 |
| PF3D7_0937500 | rifin | 0.000513242 | 2.00333E-05 | 0.039032785 |
| PF3D7_0833400 | rifin | 0.008025774 | 0.000220366 | 0.027457267 |
| PF3D7_1479400 | rifin | 0.001103529 | 2.00333E-05 | 0.018153809 |
| PF3D7_1373000 | rifin | 0.003859388 | 4.00665E-05 | 0.01038157 |
| PF3D7_1373300 | rifin | 0.021379542 | 0.000200333 | 0.009370292 |
| PF3D7_0324800 | rifin | 7.82308E-05 | 0 | 0 |
| PF3D7_1254400 | rifin | 1.06678E-05 | 0 | 0 |
| PF3D7_0300700 | rifin | 5.80805E-05 | 0 | 0 |
| PF3D7_1040500 | rifin | 7.58602E-05 | 0 | 0 |
| PF3D7_0617700 | rifin | 9.48253E-06 | 0 | 0 |
| PF3D7_1300200 | rifin | 8.65281E-05 | 0 | 0 |
| PF3D7_0901100 | rifin | 4.74126E-06 | 0 | 0 |
| PF3D7_1479600 | rifin | 1.06678E-05 | 0 | 0 |
| PF3D7_1373100 | rifin | 0.010927427 | 0 | 0 |
| PF3D7_0222600 | rifin | 0.000183724 | 0 | 0 |
| PF3D7_1100300 | rifin | 0.000158832 | 0 | 0 |
| PF3D7_1150300 | rifin | 1.42238E-05 | 0 | 0 |
| PF3D7_0701800 | rifin | 2.48916E-05 | 0 | 0 |
| PF3D7_1150200 | rifin | 4.14861E-05 | 0 | 0 |
| PF3D7_0713000 | rifin | 0.00024299 | 0 | 0 |
| PF3D7_1400400 | rifin | 5.92658E-06 | 0 | 0 |
| PF3D7_1100400 | rifin | 5.92658E-05 | 0 | 0 |
| PF3D7_1479700 | rifin | 7.23043E-05 | 0 | 0 |
| PF3D7_0114700 | rifin | 0.000293958 | 0 | 0 |
| PF3D7_1000400 | rifin | 2.01504E-05 | 0 | 0 |
| PF3D7_1300400 | rifin | 0.000112605 | 0 | 0 |
| PF3D7_0200700 | rifin | 2.60769E-05 | 0 | 0 |
| PF3D7_0425900 | rifin | 1.30385E-05 | 0 | 0 |
| PF3D7_0900200 | rifin | 0.000286846 | 0 | 0 |
| PF3D7_1400200 | rifin | 0.001059672 | 0 | 0 |
| PF3D7_1372700 | rifin | 0.000101937 | 0 | 0 |
| PF3D7_0200600 | rifin | 4.03007E-05 | 0 | 0 |
| PF3D7_0100400 | rifin | 3.91154E-05 | 0 | 0 |
| PF3D7_0833100 | rifin | 0.000164759 | 0 | 0 |

**Extended Data Table 7.**
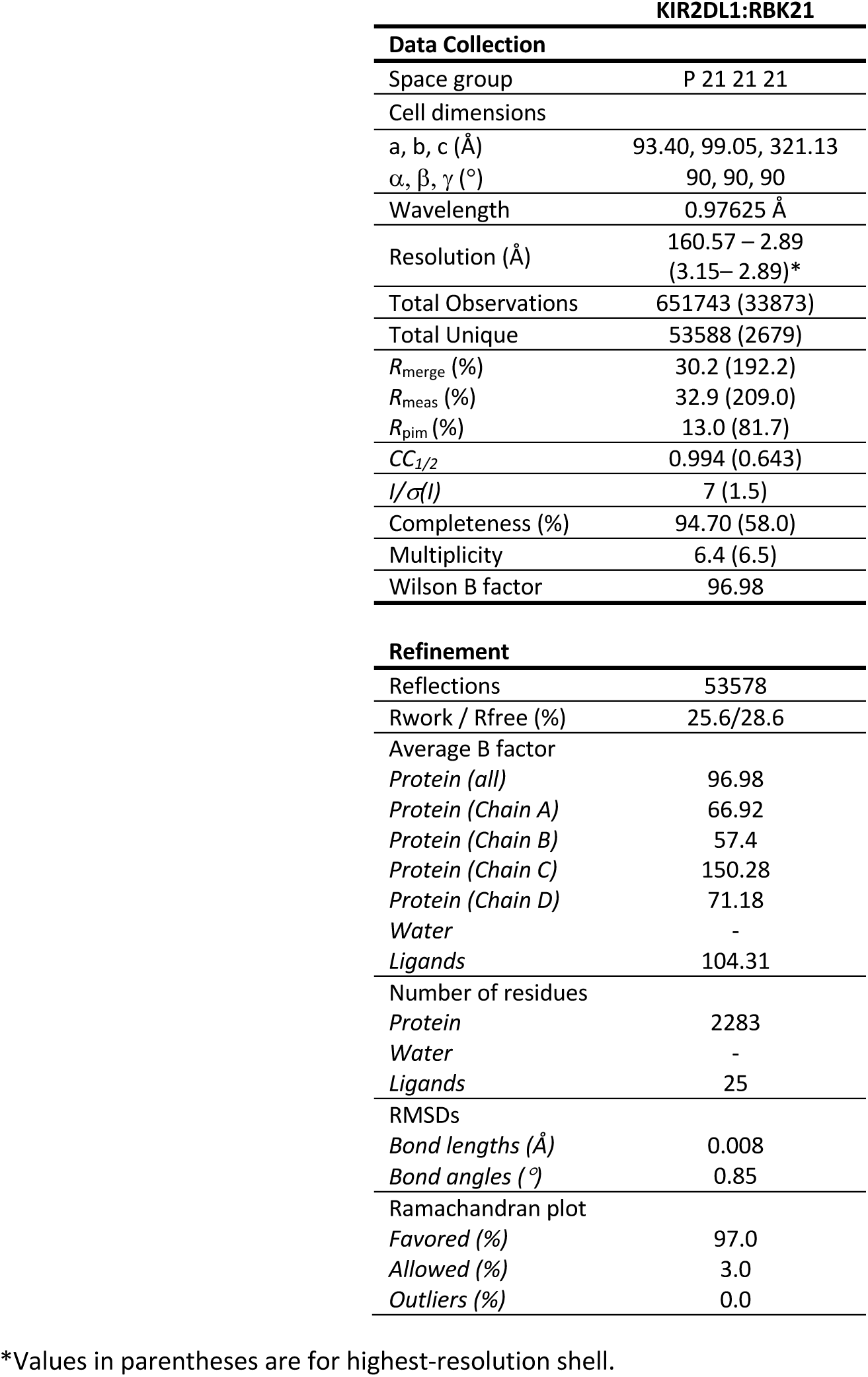
Crystallographic data collection and refinement statistics.

| KIR2DL1:RBK21 |  |
| --- | --- |
| **Data Collection** |  |
| Space group | P 21 21 21 |
| Cell dimensions |  |
| a, b, c (Å) | 93.40, 99.05, 321.13 |
| $\alpha$, $\beta$, $\gamma$ (°) | 90, 90, 90 |
| Wavelength | 0.97625 Å |
| Resolution (Å) | 160.57 – 2.89<br>(3.15– 2.89)* |
| Total Observations | 651743 (33873) |
| Total Unique | 53588 (2679) |
| $R_{\text{merge}}$ (%) | 30.2 (192.2) |
| $R_{\text{meas}}$ (%) | 32.9 (209.0) |
| $R_{\text{pim}}$ (%) | 13.0 (81.7) |
| $CC_{1/2}$ | 0.994 (0.643) |
| $I/\sigma(I)$ | 7 (1.5) |
| Completeness (%) | 94.70 (58.0) |
| Multiplicity | 6.4 (6.5) |
| Wilson B factor | 96.98 |
| **Refinement** |  |
| Reflections | 53578 |
| Rwork / Rfree (%) | 25.6/28.6 |
| Average B factor |  |
| Protein (all) | 96.98 |
| Protein (Chain A) | 66.92 |
| Protein (Chain B) | 57.4 |
| Protein (Chain C) | 150.28 |
| Protein (Chain D) | 71.18 |
| Water | - |
| Ligands | 104.31 |
| Number of residues |  |
| Protein | 2283 |
| Water | - |
| Ligands | 25 |
| RMSDs |  |
| Bond lengths (Å) | 0.008 |
| Bond angles (°) | 0.85 |
| Ramachandran plot |  |
| Favored (%) | 97.0 |
| Allowed (%) | 3.0 |
| Outliers (%) | 0.0 |

\*Values in parentheses are for highest-resolution shell.

**Extended Data Figure 8:** Table of contacts between RBK21 and KIR2DL1.

| Chain (RBK21) | Residue | Group | Chain (KIR2DL1) | Residue | Group | Interaction |
| --- | --- | --- | --- | --- | --- | --- |
| A | S221 | Side chain | B | T69 | Side chain | H bond |
| C |  |  | D | N67 | Main chain | H bond |
| A | S222 | Side chain | B | N67 | Glycan | H bond |
| A | L224 | Side chain | B | L59 | Side chain | Hydrophobic |
|  |  |  |  | H61 |  |  |
|  |  |  |  | T69 |  |  |
|  |  |  |  | Y101 |  |  |
|  |  |  |  | V111 |  |  |
| A | T225 | Side chain | B | N67 | Glycan | H bond |
| A | M239 | Side chain | B | Y109 | Side chain | Hydrophobic |
| A | F242 | Side chain | B | L59 | Side chain | Hydrophobic |
| A |  | Side chain | B | T69 |  | Hydrophobic |
| A |  | Side chain | B | R71 |  | Hydrophobic |
| A |  | Main chain | B | R71 |  | H bond |
| C |  | Main chain | D | H57 |  | H bond |
| A | F243 | Side chain | B | L59 | Side chain | Hydrophobic |
|  |  |  |  | T69 |  |  |
| A | A245 | Main chain | B | R71 | Side chain | H bond |
| A | V257 | Main chain | B | R71 | Main chain | H bond |
| A | S258 | Side chain | B | D68 | Side chain | H bond |
| A | M259 | Main chain | B | T69 | Main chain | H bond |
| A | N260 | Side chain | B | D68 | Side chain | H bond |
|  |  | Main chain |  | D68 | Side chain | H bond |

## Notes

### Competing Interest Statement

The authors have declared no competing interest.

